# Self-organized traveling waves in a synthetic multicellular reaction-diffusion system

**DOI:** 10.64898/2026.06.24.734269

**Authors:** Amit Landge, Luciano Marcon, Gary Soh, Lennart Lohrmann, Stefan Volkwein, Patrick Müller

## Abstract

Synthetic gene circuits provide experimentally tractable systems for dissecting how genetic feedback and diffusible signals generate multicellular patterns. However, building multi-component circuits whose behavior can be quantitatively linked to module-level measurements, diffusion, and spatial boundary conditions remains challenging. Here, we designed and engineered a bacterial patterning system in which positive feedback, delayed negative feedback, and two orthogonal quorum-sensing signals are integrated in *Escherichia coli*. We first implemented and characterized the feedback modules separately, measured the effective diffusion of the signals in the experimental setup, and used these data to parameterize a mathematical model. In quasi-2D bacterial lawns, the complete circuit generated self-organized spatiotemporal dynamics consisting of an sfGFP activation front followed by successive mCherry propagating pulses/traveling waves. Model-guided perturbations showed that lawn size, lawn position relative to the domain boundary, and signal degradation modulate the timing, amplitude, wavelength, and directionality of these patterns. Our work establishes a modular synthetic multicellular reaction-diffusion system in which circuit architecture, signal diffusion, and boundary-mediated signal exchange can be experimentally connected to emergent patterning dynamics.

## Introduction

Multicellular patterns emerge when local genetic feedback is coupled to communication between cells. In natural systems, diffusible signals and feedback control diverse patterning processes, including gradient formation, propagating waves, and spatially periodic patterns^1–15^. Which output emerges depends on network topology, feedback strength, feedback delays, signal degradation, diffusion range, and the geometry in which the system operates^8,11,12,16–24^. Because these factors are tightly intertwined in natural tissues, synthetic genetic circuits provide a complementary strategy to implement defined network architectures, measure their components, and experimentally perturb the physical context in which collective dynamics emerge^9,25–29^.

Synthetic genetic circuits have increasingly been used to engineer and analyze multicellular pattern formation^9,10,19,25,27,30–38^. Compared with endogenous developmental networks, these systems can be programmable, tunable, and modular, making them well suited for testing how circuit architecture and intercellular communication shape collective behavior^27,36^. For example, activator-inhibitor circuits have been implemented to generate stochastic spatial patterns of gene expression in bacteria^10^ and mammalian cell culture^9^, and bacterial colonies have been engineered to form periodic growth patterns based on local activation and long-range inhibition^30^. Quorum-sensing circuits have also been used to couple genetic clocks and oscillator networks across bacterial populations^32,34,39^, including systems in which synchronized genetic oscillations or spontaneous traveling waves of gene expression emerge in spatially extended colonies^34^. Microbial consortia carrying complementary activator and repressor circuits provide another route to multicellular dynamics driven by diffusible signals^37^. In parallel, oscillator designs such as the repressilator^38^ and its later optimized variants^40^ have established principles for generating robust temporal dynamics, while recent bacterial reaction-diffusion circuits have produced stripes, spots, and traveling wave-like patterns in different parameter regimes^25^. These studies established important principles for synthetic multicellular patterning, but they also highlight the need for additional experimentally tractable multicomponent systems in which module-level characterization, measured signal diffusion, spatial boundary conditions, and model predictions are connected within one framework.

The design of such synthetic experimental platforms is inspired by natural patterning systems. Diffusion-mediated signaling waves and spatial patterns occur in several biological contexts^7,41^. For example, traveling waves of growth-factor signaling contribute to zebrafish scale growth and regeneration, and mathematical modeling suggests that inhibitory feedback may be important for these dynamics^42^. Spatially constrained stem-cell micropatterns further illustrate how geometry, edge-mediated signal loss, and endogenous signaling networks can combine to produce reproducible differentiation patterns^18,20,24^. However, because natural patterning systems contain many interacting components, it is often difficult to isolate the contributions of individual feedback loops, diffusible signals, and boundary conditions experimentally^43^. Synthetic systems therefore provide a useful complementary approach: They do not replace system-specific developmental models, but allow assumptions about feedback, diffusion, and boundary-mediated signal exchange to be physically implemented and perturbed in a controlled multicellular setting.

In this study, we designed and engineered a bacterial multicellular patterning system that integrates positive feedback, delayed negative feedback, and two orthogonal diffusible quorum-sensing signals in *Escherichia coli* (*E. coli*). We first implemented and characterized the feedback modules separately, measured effective signal diffusion in the experimental setup, and used these data to parameterize a mathematical model of the complete circuit. In quasi-2D bacterial lawns, the complete circuit generated self-organized spatiotemporal dynamics consisting of an sfGFP activation front followed by successive mCherry propagating pulses/traveling waves within the 24 h imaging window. Model-guided perturbations of lawn size, lawn position relative to the domain boundary, and signal degradation showed that geometry and boundary-mediated signal exchange modulate the timing, amplitude, wavelength, and directionality of the observed patterns. Thus, our work establishes a modular bacterial reaction-diffusion platform in which circuit architecture, diffusible signaling, and physical boundary conditions can be quantitatively linked to emergent multicellular patterning dynamics.

## Results

### Design of a synthetic genetic system for pattern formation

To design a multicellular patterning system capable of generating traveling-wave-like dynamics, we considered minimal activator-inhibitor topologies that combine diffusible signals with delayed feedback^12,44,45^. Our design was guided by three principles: First, diffusible cell-cell signaling provides spatial coupling between cells in a multicellular population^33,35,46^; second, positive feedback amplifies local signal production and reporter activation; and third, delayed negative feedback can generate pulsatile or oscillatory dynamics (Fig. 1a). We implemented this architecture in *E. coli* using well-characterized acyl-homoserine lactone (AHL) quorum-sensing components^27^. Starting from a simple activator-inhibitor topology (Fig. 1a), we expanded the design into experimentally implementable genetic feedback modules modules^12^ (Fig. 1b). This modular architecture allowed us to construct and characterize the positive- and negative-feedback modules separately before combining them into the complete circuit (Fig. 1b,c, Supplementary Fig. 1).

**Figure 1.**
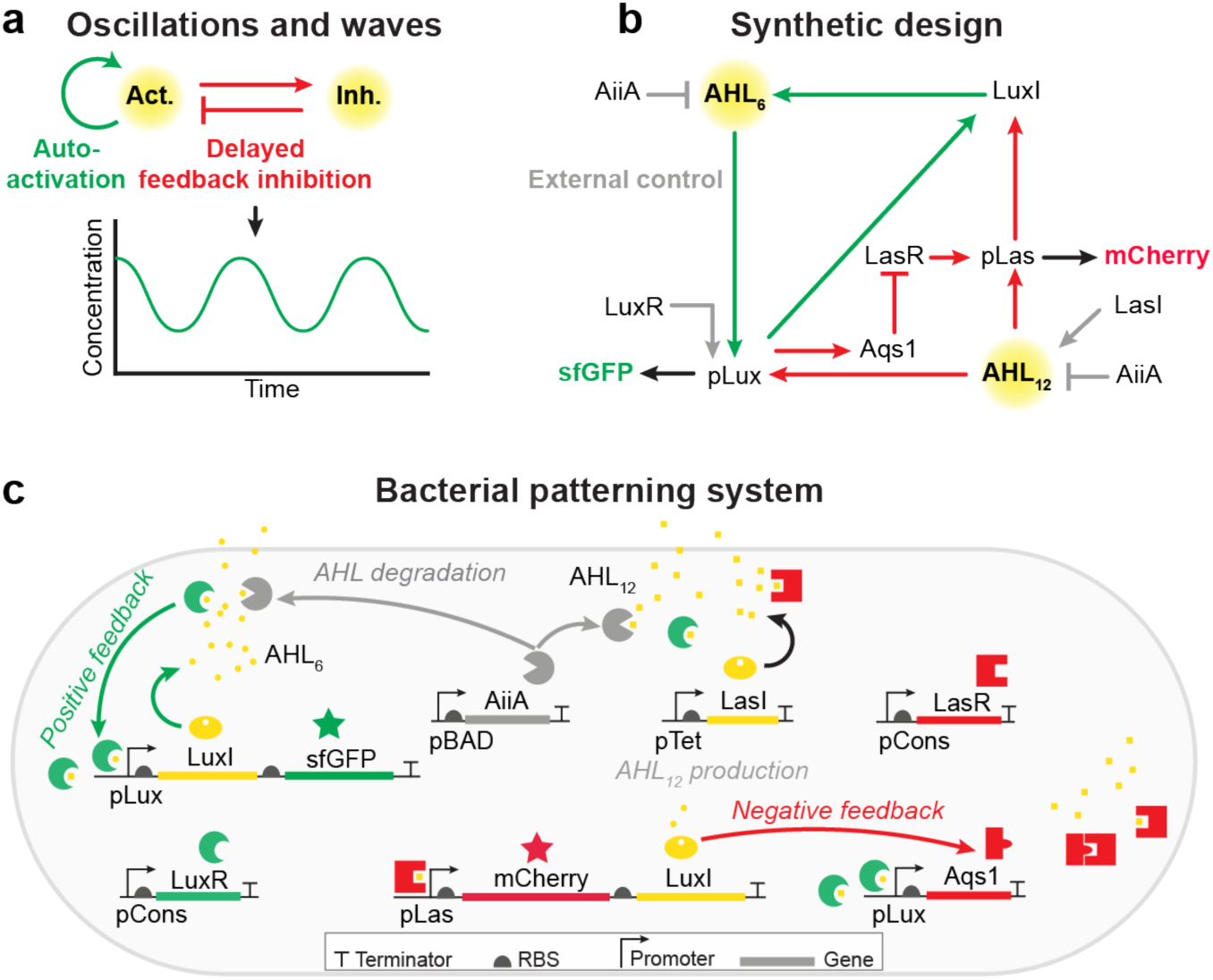
Design of a synthetic multicellular system with genetic feedback and diffusible signals. **(a)** Two-component systems with positive and delayed negative feedback can generate oscillations and waves. **(b)** Network design to implement patterning with a positive (green) and a negative feedback module (red). Diffusible components are indicated by a yellow halo. The external controls in the circuit are shown in gray. AHL6: N-(3-Oxohexanoyl)-L-homoserine lactone ligand, AHL12: N-(3-Oxododecanoyl)-L-homoserine lactone ligand, AiiA: Lactonase, LuxI: AHL6 synthase, LasI: AHL12 synthase, LuxR and LasR: ligand-binding proteins to activate *pLux* and *pLas* promoters respectively, Aqs1: anti quorum sensing protein 1, sfGFP and mCherry: fluorescent reporters to monitor dynamic circuit output. **(c)** Schematic of a bacterial cell engineered to execute the synthetic network with genetic feedbacks. AHL6 signal provides positive feedback to drive more AHL6 synthesis. The *pLas* system promotes expression of Aqs1, which provides negative feedback for *pLas* activity. The AHLs are degraded by Lactonase (*AiiA*) driven under an arabinose-inducible pBAD promoter. AHL12 production by LasI is under the control of the anhydrotetracycline-inducible pTet system. pCons: constitutively active promoter for *LasR* and *LuxR* expression, RBS: ribosome-binding-site.

In our circuit design, the positive feedback module was designed with AHL6 (N-3-(oxohexanoyl)-L-homoserine lactone) as the diffusible signal. AHL6 is a well-characterized signaling molecule from *Vibrio fischeri* that binds to its receptor LuxR to activate the *pLux* promoter^47^. We implemented an AHL6-based positive feedback module by coupling the expression of the AHL6 synthase LuxI^48^ with the *pLux* promoter (Fig. 1b,c). To provide a dynamic readout of the positive feedback module output, we placed a destabilized superfolder GFP (*sfGFP-ssrA*)^49,50^ reporter under the control of the *pLux* promoter. To externally tune the output of the circuit, we added an arabinose-inducible AHL6-degrading Lactonase (*AiiA*) from *Bacillus thuringiensis*^51^ to the system. Thus, our positive feedback module design uses AHL6 to trigger the expression of LuxI, which in turn increases AHL6 production whose levels can be regulated by the addition of arabinose. The green fluorescent signal of sfGFP serves as a readout of the module’s activity.

We built an orthogonal negative feedback module using another diffusible signal, AHL12 (N-(3-Oxododecanoyl)-L-homoserine lactone) from *Pseudomonas aeruginosa*^52^ (Fig. 1c). We made AHL12 production tunable by placing the AHL12 synthase LasI under the control of the anhydrotetracycline-inducible *pTet* promoter. AHL12 exerts its function by binding to the receptor LasR to activate the *pLas* promoter, but it can also promiscuously bind to LuxR to activate the *pLux* promoter. To implement strong negative feedback, we took advantage of this promiscuity, coupled AHL6 synthase expression to the *pLas* promoter, and placed the LasR antagonist Aqs1 (anti-quorum sensing protein 1)^53^ under control of the *pLux* promoter, which can be both activated by AHL12-LuxR and AHL6-LuxR complexes. As in the case of the positive feedback module, LuxR and LasR were designed to be expressed from constitutive promoters. Finally, a destabilized mCherry-ssrA reporter was used in the negative feedback module as the *pLas* activity readout. In this design, AHL12 produced by LasI should activate *Aqs1* expression via LuxR binding, resulting in the quenching of LasR and decreased mCherry fluorescent readout.

This design represents the foundation of our mixed positive and negative feedback system, which our mathematical analyses predict to be sufficient to generate oscillatory dynamics and traveling wave patterns (Supplementary Note).

### Characterization of the positive feedback module

To test the functionality of the positive and negative feedback modules, we constructed and characterized each of them separately. For the positive feedback module (Fig. 2a), we generated four different plasmids – pAL101, pAL102, pAL103, and pAL104 (Supplementary Fig. 1) – with varying expression levels of the key mediator LuxR to screen for optimal circuit output and dynamic range. The plasmids pAL101 and pAL103 contain different *pLux* promoter sequences, with pAL103 having a more stringent version^54^. The plasmids pAL102 and pAL104 were generated by deletion of the *LuxR* cassette from the pAL101 and pAL103 plasmids, respectively, to assess circuits with low LuxR levels. We tested all plasmids in the *E. coli* Marionette strain, which carries highly optimized small-molecule sensors and contains a chromosomal copy of *LuxR* under the control of a constitutive promoter^54^.

**Figure 2.**
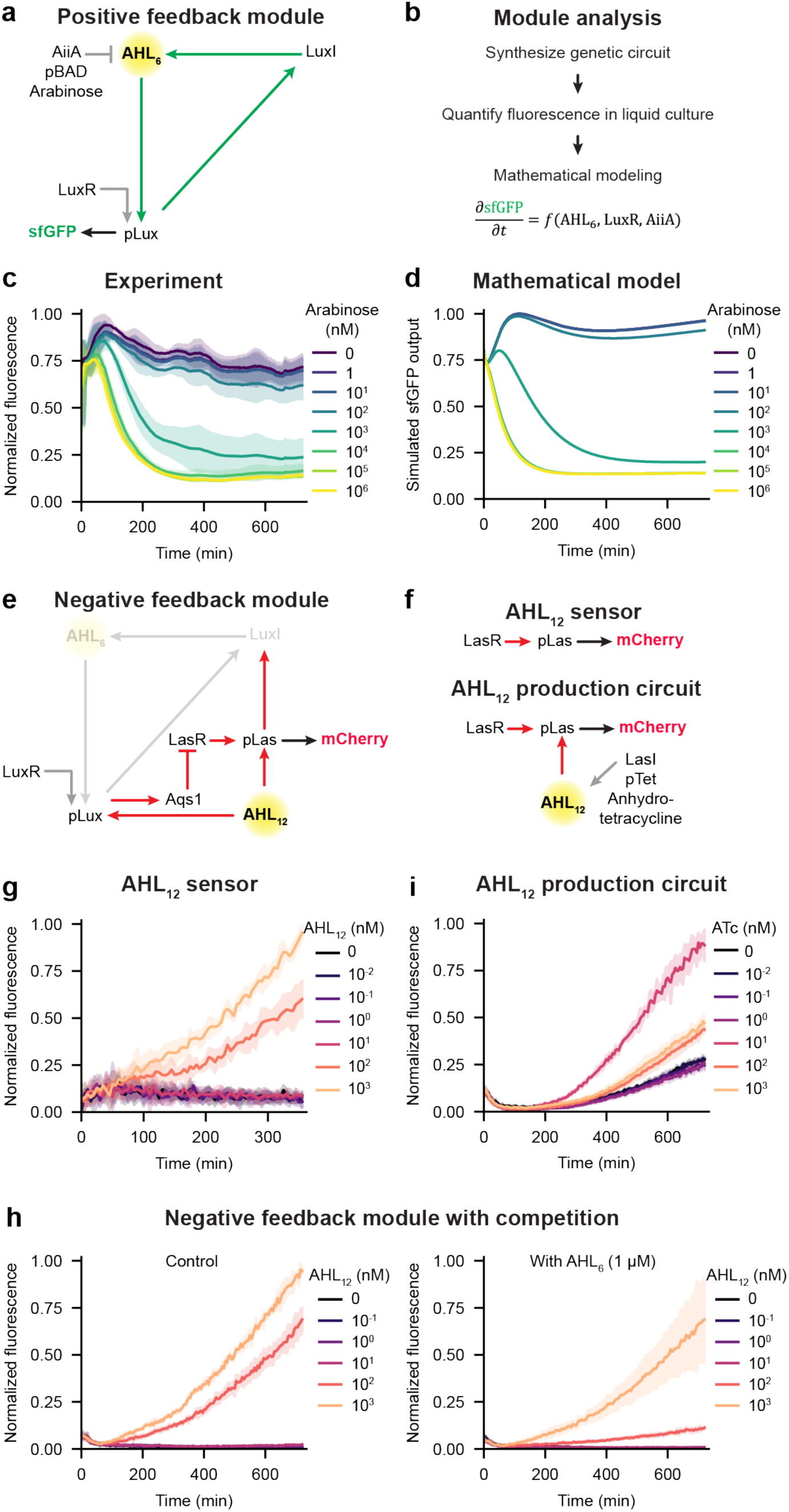
Characterization of positive and negative feedback modules. **(a)** Schematic of the positive feedback module. The AHL6 signal induces expression of LuxI (AHL6 synthase) under the *pLux* promoter. The activator protein LuxR is expressed from a constitutive promoter. AHL degradation can be triggered through arabinose-inducible expression of AiiA (Lactonase). **(b)** Strategy to characterize the modules from circuit synthesis to experimental measurements and mathematical modeling. **(c)** Quantification of sfGFP fluorescence (normalized by OD600) for the positive feedback module (plasmid pAL103) under external arabinose control. n = 3 biological replicates with 2 technical replicates each. **(d)** Mathematical simulation results for the experiment in (c). **(e)** Schematic of the negative-feedback module. **(f)** AHL12 sensor and AHL12 production submodules constitute the negative feedback module. **(g)** Quantification of mCherry fluorescence intensity (normalized by OD600) in the AHL12 sensor submodule for different AHL12 doses. n = 2 biological replicates with 2 technical replicates each. **(h)** Comparison of mCherry fluorescence intensity (normalized by OD600) in the full negative feedback module for different AHL12 doses without AHL6 (left) and with 1 µM AHL6 (right) is shown. n = 1 biological replicate with 3 technical replicates. **(i)** Quantification of mCherry fluorescence intensity (normalized by OD600) in the AHL12 production submodule for different ATc doses. n = 1 biological replicate with 3 technical replicates. Lines and error envelopes in (c), (g), (h), and (i) show mean and standard deviation.

To quantify the dynamics of the positive feedback module (Fig. 2a) in engineered *E. coli*, we first measured sfGFP intensity in homogeneous liquid culture (Fig. 2b). We observed that the strains carrying pAL101 and pAL103 maintained high levels of sfGFP output for up to 12 h in liquid culture at 37°C (Fig. 2c, Supplementary Fig. 2a-c, Supplementary Fig. 3a-c), while strains carrying the plasmids pAL102 and pAL104 showed only transiently high levels of sfGFP intensity (Supplementary Fig. 4a-d, Supplementary Fig. 5a-c), consistent with the lower *LuxR* copy number and reduced *LuxR* mRNA levels (Supplementary Fig. 4d). We designed the positive feedback module such that arabinose-inducible expression of Lactonase (*AiiA*) should be able to control the output of the circuit by modulating the degradation of AHL6. Indeed, upon arabinose treatments the mRNA levels of *AiiA* were increased (Supplementary Fig. 2d, Supplementary Fig. 4e). Moreover, *LuxI* and *sfGFP* mRNA levels correspondingly decreased for the pAL101 strain, while *LuxR* levels were not strongly affected (Supplementary Fig. 2e-g). The dose-response analysis also showed a quantitative decrease in sfGFP fluorescence intensities (Supplementary Fig. 2h, Supplementary Fig. 3d, Supplementary Fig. 4f, Supplementary Fig. 5d). The inhibitory threshold concentration of arabinose determined by fitting the data was in the range of 480 nM to 1400 nM (Supplementary Table 2), and the pAL103 circuit with the best dynamic range (Supplementary Table 2) was selected for further analysis. Parameterization of our mathematical model with these measurements closely mimicked the experimental findings (Fig. 2d, Supplementary Fig. 3e, Supplementary Note), rationalizing the use of the positive feedback model for the full patterning circuit.

### Characterization of the negative feedback module

To simplify the implementation and characterization of the negative feedback module (Fig. 2e), we further divided it into submodules: AHL12 sensor, AHL12 production, and LasR inhibition submodules (Fig. 2e-f, Supplementary Fig. 1a,c, Supplementary Fig. 6a). The AHL12 sensor plasmid was designed with a destabilized mCherry (mCherry-ssrA) expressed under the AHL12-inducible *pLas* promoter. The activator protein LasR, needed to form the AHL12-LasR complex, was expressed under a constitutive promoter (Supplementary Fig. 1a). We characterized the dose-response behavior of the sensor circuit output in terms of mCherry fluorescence with different levels of exogenous AHL12. We found a robust induction with high dynamic range, low leakiness, and an activation threshold concentration of 76 ± 4 nM (Fig. 2g, Supplementary Fig. 6b-d). Moreover, qRT-PCR analysis showed that *mCherry* expression was AHL12-specific (Supplementary Fig. 6f), and AHL12 doses greater than 1 nM induced high *mCherry* expression (Supplementary Fig. 6g), while *LasR* mRNA levels were not strongly affected (Supplementary Fig. 6h-i). Based on these results, we parameterized our mathematical model for the AHL12 sensor submodule (Supplementary Note), which closely matched the observed mCherry output dynamics and dose-response profile (Supplementary Fig. 6e).

We implemented the negative feedback on mCherry output through inhibition of the activator protein LasR mediated by the antagonist Aqs1^53^, and we hypothesized that the AHL12 threshold should be increased through Aqs1-mediated LasR inhibition. To test this idea, we generated the LasR inhibition submodule by placing *Aqs1* under the control of the *pLux* promoter, which was co-transformed with the AHL12 sensor plasmid into the Marionette strain. We first tested whether the LasR inhibition submodule, carrying both the AHL12 sensor and LasR inhibition plasmids, could yield tunable mCherry fluorescence output upon treatment with different AHL12 concentrations (Fig. 2h). The activation threshold concentration of AHL12 was 93 ± 2 nM (Supplementary Fig. 6j). This activation threshold was higher than that of the AHL12 sensor submodule (Supplementary Fig. 6d), since AHL12 in the LasR inhibition submodule can also partially activate *Aqs1* expression by promiscuity with the *pLux* promoter (Supplementary Fig. 6l). To test the competition between Aqs1 and AHL12 for the binding to LasR, we induced Aqs1 expression by adding 1 µM AHL6. As a result, the AHL12 threshold increased to 160 ± 22 nM (Fig. 2h, Supplementary Fig. 6j). Thus, Aqs1 was able to inhibit LasR and tune the output of the negative feedback module, in good agreement with the output of our mathematical model (Supplementary Fig. 6k).

In our synthetic network (Fig. 1c), the input of AHL12 needs to be dynamically controlled. To this end, we placed the AHL12 synthase *LasI* under the control of the anhydrotetracycline (ATc)-inducible *pTet* promoter^55^ to generate the AHL12 production submodule (Supplementary Fig. 1c). The corresponding plasmid was designed such that it can be maintained along with the AHL12 sensor plasmid in the Marionette strain. A dose-response assay was performed for the resulting AHL12 production submodule to quantify mCherry fluorescence with different ATc concentrations (Fig. 2i). The highest fluorescence normalized to cell density (OD600) was achieved with a 10 nM ATc treatment (Fig. 2i, Supplementary Fig. 6n). As expected, qRT-PCR analysis showed that *LasI* and *mCherry* mRNA levels increased in a dose-dependent manner upon ATc treatment (Supplementary Fig. 6o,p, Supplementary Fig. 7), which saturated at very high levels above 100 nM. These data show that basal *pLas* promoter activation in the AHL12 production strain can be robustly boosted by external control within physiological levels.

Taken together, our validation studies of positive and negative feedback modules confirmed that they function as designed and stipulated by our mathematical model.

### Measuring AHL diffusion rates using sensor strains

Spatial coupling through diffusion of signaling molecules is essential for the generation of patterns in our circuit, and the wave length of the emerging structures is dependent on the magnitude of the diffusion rates. To determine the diffusion coefficients of the AHL signaling molecules in our experimental setup, we used sensor strains that provide a quantitative, dose-dependent fluorescent readout upon AHL treatment (Fig. 3a, Fig. 2g, Supplementary Fig. 1a, Supplementary Fig. 8). We added a known amount of AHL at the center of a lawn of sensor cells to create a signal source. The gradient of AHL concentration created by diffusion from the source led to a fluorescent reporter signal gradient (Fig. 3a). Dynamic radial profiles of this fluorescence gradient were quantified at multiple time points as the AHL molecules diffused into the sensor cell lawn. Mathematical models of the experimentally validated sensor strain dynamics were used to obtain simulated radial profiles (Supplementary Note), and the AHL diffusion coefficients were determined by fitting the simulated data to the experimental profiles (Fig. 3b,c). To validate the method, we performed measurements and fits with gradients generated by fluorescein-labeled dextran molecules under the same experimental conditions and using the same fitting methods (Fig. 3d,e). We found diffusion coefficients of 120 ± 5 μm^2^/s and 67 ± 12 μm^2^/s for 3 kDa and 10 kDa dextrans, respectively (Fig. 3f, Supplementary Table 3), similar to previously reported values^56^. For AHL6 and AHL12, we determined diffusion coefficients of 313 ± 69 μm^2^/s and 320 ± 59 μm^2^/s, respectively. These measurements are in line with the AHL’s small molecular weights of 0.2-0.3 kDa and demonstrate that the signaling molecules in our system can mediate long-range spatial coupling across millimeter-wide domains.

**Figure 3.**
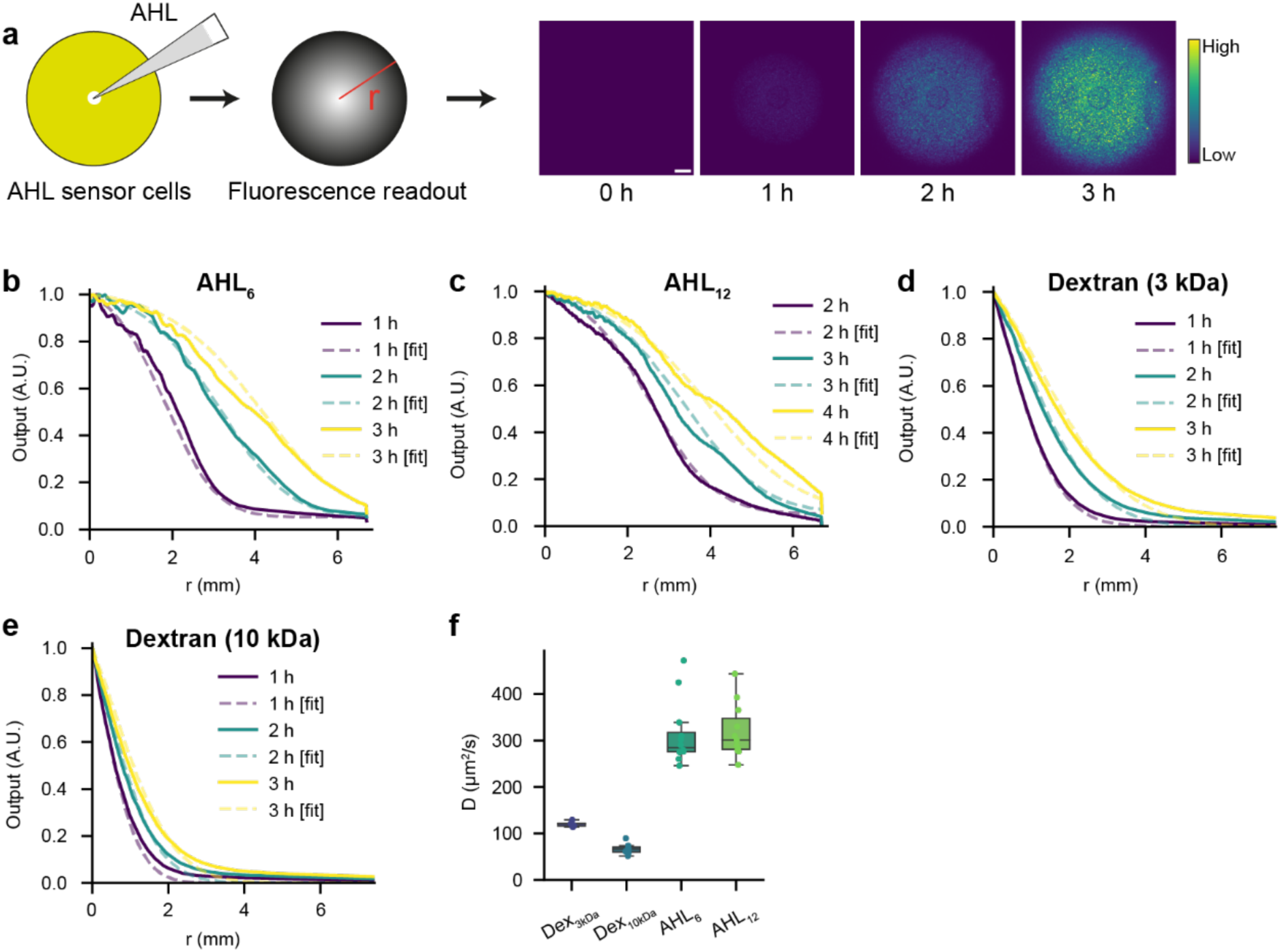
Measurement of AHL diffusion coefficients using AHL sensor strains. **(a)** Methodology to generate fluorescence gradients in an AHL sensor strain lawn by adding the inducer (AHL) to the lawn center. Gradients were quantified and fitted to simulations of a mathematical model to determine the AHL diffusion coefficient. **(b-e)** Representative fits of experimental gradients and simulated gradients for AHL6, AHL12, fluorescein-labeled 3 kDa dextran, and fluorescein-labeled 10 kDa dextran. **(f)** Boxplot showing measured diffusion coefficients. n = 12, 11, 7, and 7 for AHL6, AHL12, 3 kDa dextran, and 10 kDa dextran respectively. The center line represents the median, while the upper and lower box edges correspond to the first and third quartiles, respectively. Whiskers extend to the farthest data point within 1.5 times the interquartile range from the box edges or to the maximum data point within this range, whichever is closer.

### The synthetic multicellular system generates spontaneous traveling waves

Our design predicts the emergence of traveling waves for the full circuit comprising positive and negative feedback. To test this prediction, we generated the complete synthetic genetic network by placing the characterized submodules on two compatible plasmids (Supplementary Fig. 1d). We introduced the complete circuit into the Marionette strain to test for the emergence of spatiotemporal patterns in quasi-2D multicellular lawns with negligible thickness and a shallow cell density gradient resulting from a slightly curved agar surface (Fig. 4a). Using time-lapse imaging, we observed oscillatory dynamics and spontaneous traveling waves of mCherry fluorescence (Fig. 4b-d, Supplementary Movie 1). In this system, mCherry fluorescence initially increased uniformly for 6 h (Supplementary Movie 1). The first traveling wave was observed as the signal decayed radially outward, followed by the initiation of a second traveling wave at the lawn center at ∼10 h with a speed of about 5 μm/min. The second wave reached the edge of the lawn at ∼16 h. In contrast to the oscillatory mCherry dynamics, sfGFP showed a single peak (Fig. 4e).

**Figure 4.**
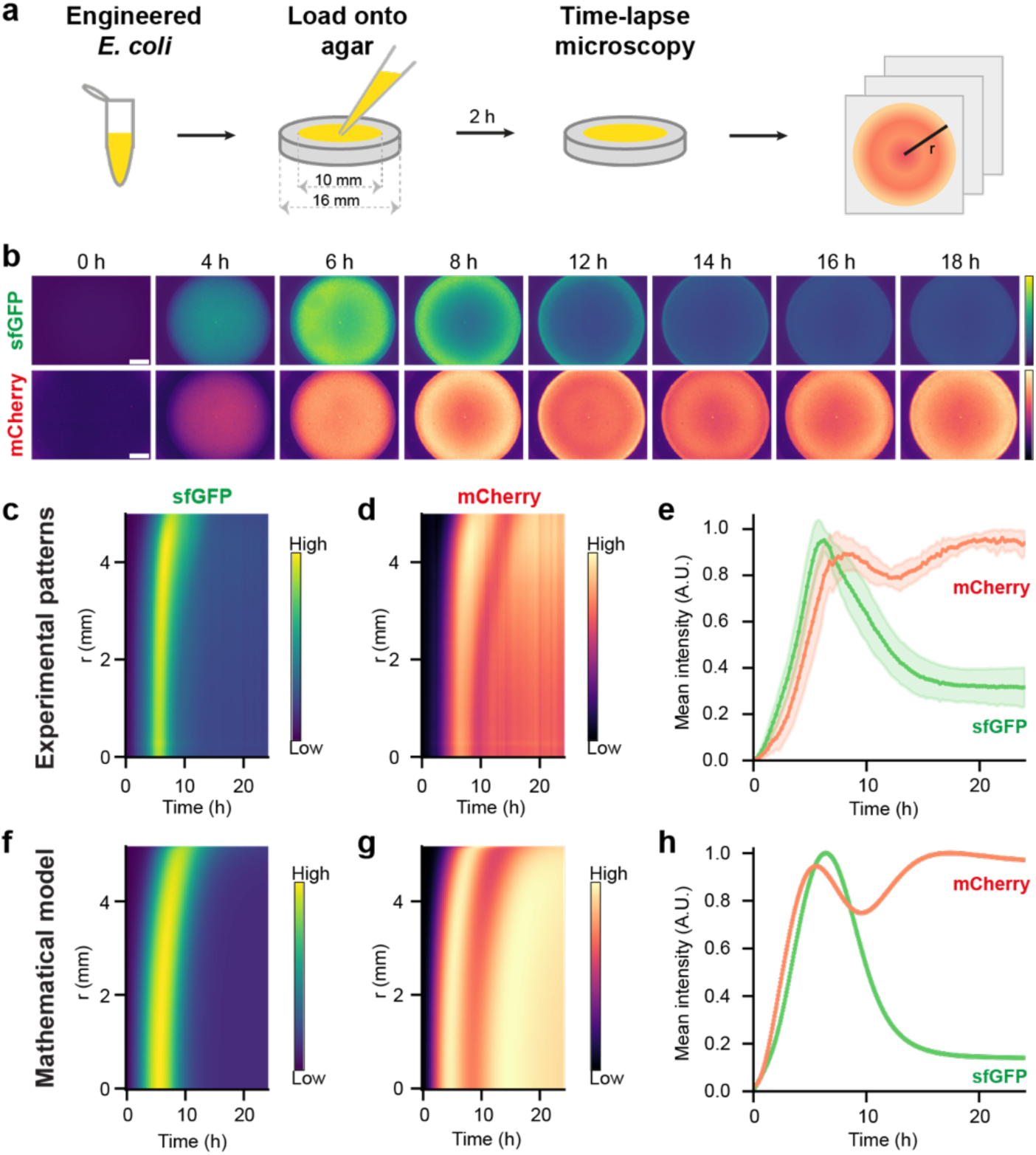
Spontaneous traveling wave patterns emerge in a lawn of engineered *E. coli*. **(a)** Workflow to assess the emergence of spatiotemporal patterns in a quasi-2D lawn of engineered *E. coli*. **(b)** Fluorescence images showing sfGFP and mCherry signals in *E. coli* lawns expressing the complete synthetic circuit (Fig. 1b,c). Scale bar: 2 mm. **(c, d)** Space-time plots showing radially averaged sfGFP (c) and mCherry (d) intensities in 15 min intervals for 24 h. **(e)** Quantification of sfGFP and mCherry mean intensities in the lawn area. n = 9 biological replicates. Error envelopes indicate standard deviation. **(f-h)** Numerical simulation results showing space-time plots of sfGFP (f) and mCherry (g) as well as mean sfGFP and mCherry outputs (h) over time.

To validate that the traveling wave patterns are specific to the synthetic circuit, we imaged the original Marionette strain (no-plasmid control) and a constitutive mCherry expression strain (control for plasmid-specific effects) using the same experimental setup (Supplementary Fig. 9, Supplementary Movie 2). The no-plasmid control showed constant and low auto-fluorescence signals for both sfGFP and mCherry, and the constitutive mCherry expression strain did not show oscillatory or traveling wave behaviors (Supplementary Fig. 9, Supplementary Movie 2). Parameterization of our mathematical model with our measured diffusion coefficients, induction thresholds, rate constants as well as initial and boundary conditions well recapitulated the emergence of patterns (Fig. 4f-h, Supplementary Note). The model showed that the patterning circuit can generate traveling waves with a shallow initial cell density gradient, whereas the circuit produces temporal oscillations with uniform initial cell density (Supplementary Fig. 10a-d, Supplementary Movie 3). Interestingly, the positive feedback module alone can also generate a single wave of sfGFP expression in this spatial setup. However, the final pattern is substantially more noisy compared to the full circuit (Supplementary Fig. 10e-j), and complex patterns – such as the pulsatile traveling waves of mCherry expression – cannot be generated.

Together, these results highlight the role of the interplay between integrated positive and negative feedback loops in reaction-diffusion systems to generate complex and regular patterns. The experimental observations combined with our mathematical analysis suggest the following intuitive model for emergence of traveling wave patterns in our system: The initial radial symmetry and cell density gradient cause rapid AHL6 synthesis and a radial wave of sfGFP output via the positive feedback circuit. In parallel, the negative feedback circuit drives AHL12 production and emergence of an mCherry wave. Next, the dynamic spatial distribution of AHL6 and AHL12 via diffusion leads to negative feedback activation in the cell lawn leading to traveling waves of mCherry expression. At ∼12 h, two radial waves of mCherry can be observed. Later, the sfGFP distribution becomes homogeneous, possibly due to homogenous AHL6 distribution via diffusion into the agar and degradation. Finally, mCherry wave propagation is halted giving a ring of mCherry at the lawn edge (Figure 4b).

### Perturbation analysis reveals the role of geometry for pattern formation

The novel patterning circuit allows us to address fundamental questions that could previously not be directly investigated in complex natural biological patterning systems^7,18,20,41,42^. Specifically, our mathematical model makes three major predictions regarding the interaction of the domain geometry and environment with the emerging patterning collective. First, the traveling wave dynamics should be dependent on the initial bacterial lawn size; smaller lawns should exhibit wider waves since the circuit inhibitor can diffuse into a larger sink domain, whereas in bigger lawns the second mCherry wave should appear earlier (Fig. 5b-e). Second, the direction of wave propagation should be modulated by the position of the domain; placing the lawn off-center closer to a boundary should reflect the diffusing signal molecules and therefore lead to anisotropic wave propagation (Fig. 5j-k,n-o). Third, the emergence and characteristic length of traveling waves should be controllable by modulating AHL signal levels, and strongly increasing the degradation of AHL signals should abrogate diffusion-dependent patterning (Supplementary Fig. 11d).

**Figure 5.**
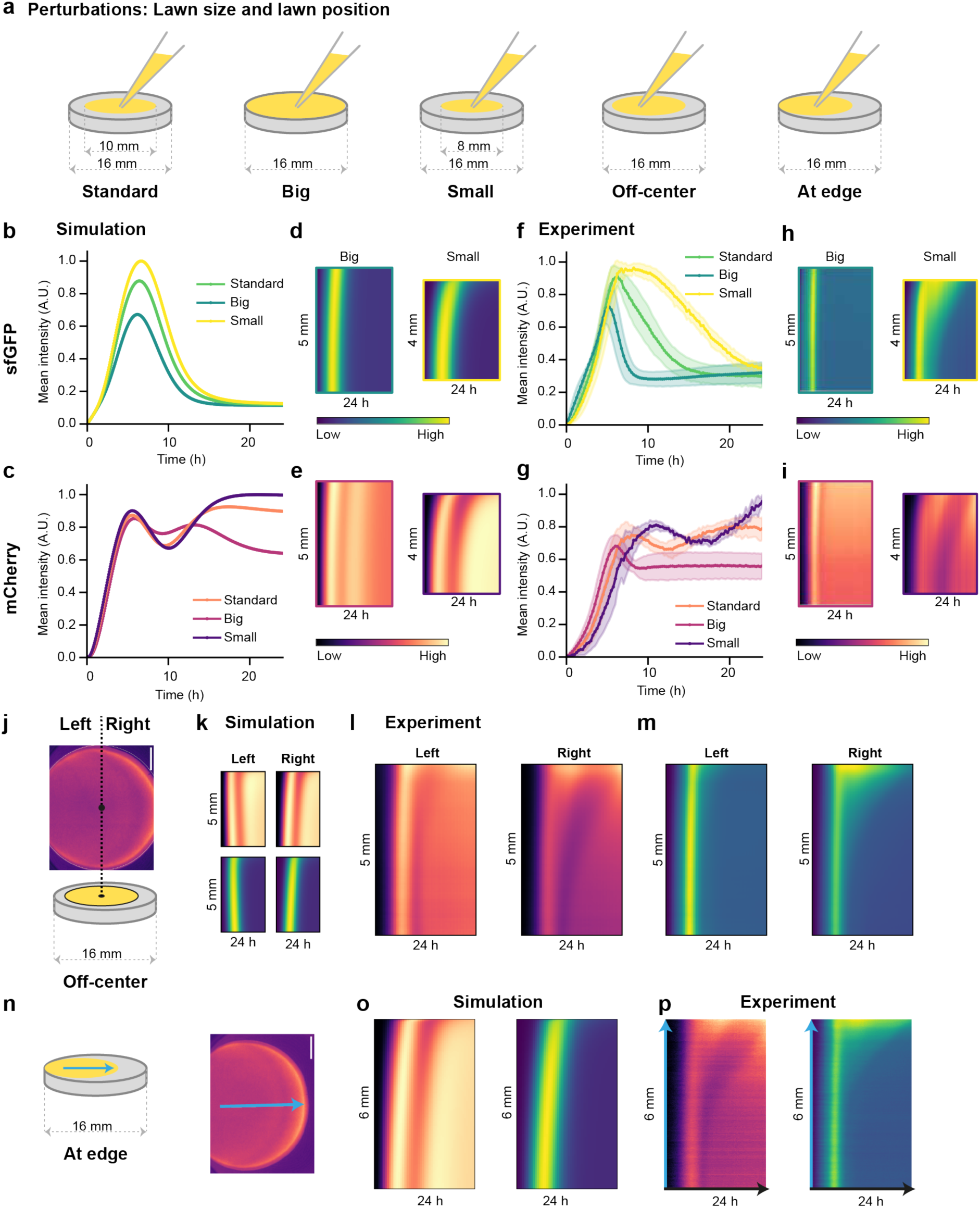
Perturbation analysis reveals the influence of domain geometry and boundary effects on traveling wave patterns. **(a)** Schematic of domain size and shape alterations as perturbations to assess boundary effects on pattern formation. **(b,c)** Simulated sfGFP (b) and mCherry (c) mean intensities for the three lawn sizes. **(d,e)** Simulated space-time plots of sfGFP (d) and mCherry (e) for the big and small lawn sizes. **(f,g)** Mean intensities of sfGFP (f) and mCherry (g) in the lawn area for the different lawn sizes. n = 6 biological replicates for big and small lawns. Error envelopes indicate standard deviation. The data for standard lawn size is the same as in Fig. 4e. **(h,i)** Representative space-time plots of sfGFP (h) and mCherry (i) fluorescence for big and small lawn sizes. For the big lawns, data until 5 mm are shown. n = 6 biological replicates. **(j)** Illustration of the quantification method to analyze the effect of initial conditions (off-center) on the direction of wave propagation and propagation anisotropy. A representative image of mCherry fluorescence is shown. The lawn is assigned left and right sides based on the shift of the lawn center. Radial profiles of fluorescence intensity are quantified in the semicircular region on each side. Scale bar: 2 mm. **(k)** Simulated space-time plots of mCherry and sfGFP for the off-center lawn condition. **(l,m)** Representative space-time plots of mCherry (l) and sfGFP (m) intensities on the left and right sides of the off-center lawn. n = 6 biological replicates. **(n)** Illustration showing the method to measure space-time profile for the at-edge lawn position. A representative mCherry fluorescence image is shown, and the blue arrow indicates the region used to generate space-time plots shown in (p). **(o)** Simulated space-time profiles of mCherry and sfGFP fluorescence for the at-edge lawn condition. **(p)** Space-time plots of mCherry and sfGFP fluorescence for the at-edge lawn condition. n = 6 biological replicates.

To test the first prediction of our mathematical model (Fig. 5b-e, Supplementary Movie 4), we varied the initial lawn size in our time-lapse imaging experiments (Fig. 5a). In the standard experimental setup, we loaded 20 µl of the cell suspension onto LB agar, creating a lawn of ∼10 mm diameter. We generated a small lawn of ∼8 mm diameter by loading 10 µl of cell mix, and a large lawn covering the entire well (∼16 mm diameter) by loading 40 µl of the cell suspension. We hypothesized that the diffusion of AHL molecules into the free agar not covered by the cell lawn may regulate the traveling wave dynamics. Particularly, smaller lawns have a larger cell-free agar area around them, acting as a sink. AHLs can diffuse into this area from the lawn boundary, and smaller lawns may therefore show slower traveling wave dynamics, whereas larger lawns retain almost all of the produced AHLs inside and may display faster dynamics. Indeed, we observed that the big lawn showed faster appearance and decay of sfGFP signal and earlier emergence of the second traveling wave of mCherry signal, albeit with a much lower amplitude (Fig. 5f-i, Supplementary Movie 5). Conversely, the small lawn showed slower and delayed traveling wave dynamics. The sfGFP signal decayed much slower, and the second wave of mCherry signal was delayed by ∼4 h. Quantification of mean fluorescence intensities over the lawn area further supported delayed wave dynamics in the smaller lawns (Fig. 5f,g). Therefore, the experimental findings well recapitulated our mathematical predictions (Fig. 5b-i, Supplementary Movie 4, Supplementary Movie 5, Supplementary Note).

To test the second prediction (Fig. 5a,j-k,n-o, Supplementary Movie 6), we varied the lawn position relative to the domain boundary. We observed radially isotropic wave propagation in the standard experimental setup. We asked whether the wave propagation can become anisotropic in response to varying the position of the lawn with respect to the domain boundary (Fig. 5a, Supplementary Movie 6). We tested off-center and at-edge lawn positions generated by loading 20 µl of the cell suspension at a slightly off-center position in the well and closer to the edge of the well, respectively. For the off-center lawn position, initially the sfGFP and mCherry signals appeared starting from the lawn center; however, the subsequent wave propagation was no longer radially isotropic (Fig. 5l,m, Supplementary Movie 7). We observed that the wave’s initial center no longer coincided with the lawn center, but it was shifted closer to the well boundary (left side in Fig. 5j). The wave propagation anisotropy was even more pronounced when the lawn was positioned at the edge of the well (Fig. 5n-p, Supplementary Movie 7). These results indicate that the directional loss of diffusible signals (AHLs) from the cell lawn into the cell-free agar as well as reflective boundary conditions determine the direction of the traveling waves.

To test the third prediction of our mathematical model (Supplementary Fig. 11a-d), we treated the cell lawns with arabinose to induce Lactonase expression and subsequent AHL degradation (Fig. 1). We perturbed the system with three different arabinose treatments: 1 nM, 1 µM and 1 mM. Since the threshold concentration of arabinose is in the micromolar range (Supplementary Fig. 3d, Supplementary Table 2), 1 nM treatment did not show a strong effect on the traveling waves compared to controls (Supplementary Fig. 11e-g, Supplementary Movie 8). Treatment with 1 µM arabinose reduced the sfGFP signal peak, but mCherry traveling waves were still observed (Supplementary Fig. 11e-g, Supplementary Movie 8). Strikingly, 1 mM arabinose treatment reduced the sfGFP signal almost completely, and a single peak of mCherry signal was observed (Supplementary Fig. 11e-g). Paradoxically, the mCherry peak signal was increased ∼5-fold compared to the control condition (Supplementary Fig. 11f), even though in our network both AHL6 and AHL12-dependent inputs on mCherry expression should be decreased by arabinose-dependent Lactonase expression (Fig. 1b). Our model explains these findings by a complete shutdown of *pLux* activity due to AHL6 degradation, which reduces the Aqs1 concentration to such low levels that LasR inhibition is negligible. In addition, our mathematical model predicted a 10-fold lower rate of AHL12 degradation compared to the rate of AHL6 degradation to explain the observed high mCherry levels. In agreement with this prediction, previous studies have reported that Lactonase shows slower hydrolysis rates for AHL substrates with longer acyl chains^57^. The decrease in sfGFP levels with modulated characteristic wave length and a single mCherry peak upon 1 mM arabinose treatment well recapitulated the predictions of our mathematical model (Supplementary Fig. 11f-h), supporting the role of diffusible AHL molecules as drivers of spatial coupling and traveling wave generation.

Collectively, these results validate the predictions made by our mathematical model and demonstrate that lawn size, lawn position relative to the domain boundary, and AHL signal levels are key determinants of traveling wave generation, direction, and propagation.

## Discussion

Engineering self-organizing multicellular patterning systems with controllable behavior has been a long-standing challenge in synthetic biology^26–29,31,36,58–62^. Here, we advanced the capabilities and framework for synthetic multicellular pattern formation by generating a novel multicomponent genetic circuit with feedback networks and diffusible signaling molecules. We used a modular circuit design framework for the positive and negative feedback modules. The modular design allowed us to characterize the system parts separately before building the complete synthetic circuit. Furthermore, we used quantitative experimental data to parametrize our mathematical models and predict key dynamical aspects of the experimental system. Finally, we combined the modules into a complete synthetic circuit that generated traveling waves of gene expression in a multicellular lawn of engineered *E. coli*. Using mathematical modeling and perturbation analyses, we showed that the traveling waves arise due to the interplay of feedback circuits, initial conditions, and boundary conditions. Thus, our system provides a controlled synthetic platform for analyzing how genetic feedback and physical constraints together shape multicellular patterning dynamics.

The role of negative feedbacks with delay has been previously explored in biological oscillators^7,38^. Our novel synthetic circuit experimentally confirms that delayed negative feedback is essential to achieve traveling wave patterns. In our circuit, the delayed negative feedback requires multiple intermediate steps of transcription and translation, and the oscillation period is in the range of 8-10 h leading to two oscillations during the 24-hour experimental time. In addition to delayed negative feedback, we demonstrate that boundary conditions are crucial to generate radially symmetric patterns. Previous studies in stem cell micropattern systems have suggested that loss of signaling molecules from the edge of the micropattern colony can give rise to radially symmetric patterns of gene expression^18,20^. In our system, the signaling molecules diffuse into the cell-free area of the agar from the edge of the bacterial lawn, and we can directly control the patterning outcome by modulating these boundary conditions. When the lawn center was positioned in the center of the well, we observed radially isotropic traveling wave patterns. These traveling waves became anisotropic when the center of the lawn was shifted closer to the well boundary. Thus, the boundary conditions of the patterning system provide an additional mechanism to control the directionality of the traveling waves^18,63,64^.

In our experimental setup, the system reaches a steady state after 24 hours, likely due to nutrient limitations and loss of diffusible signals. Using a rich nutrient medium or external supply of nutrients may uncover long-term circuit dynamics (Supplementary Note). Furthermore, our circuit offers numerous extensions useful for synthetic biology applications. We avoided effects of cell movement by embedding the bacterial cells in a gel of low-melting-temperature agarose, but in the context of collective multicellular behavior the role of cell motility on dynamic traveling waves of gene expression may provide new insights into emergent properties of cell collectives in the future. A recently reported bacterial reaction-diffusion circuit generated stripes, spots, and traveling wave-like patterns using a three-node topology with two diffusible nodes and one non-diffusible node^25^. Our circuit differs in architecture and uses two diffusible AHL signals with similar effective diffusion coefficients under our experimental conditions. Further modifications of our system may therefore allow exploration of additional dynamical regimes, including spatially periodic patterns, and may help test how network topology and effective diffusivity influence patterning outcomes. More broadly, replacing fluorescent reporters with outputs that alter growth, adhesion, motility, or morphogenesis could extend this platform from pattern visualization toward programmable multicellular behavior (Supplementary Note).

## Acknowledgments

We thank Daniel Čapek, Erik Kehr, Anna Kögler, Milka Kostic, and Hernán Morales-Navarrete for feedback and discussion. This project has received funding from the European Research Council (ERC) under the European Union’s Horizon 2020 research and innovation program (grant agreement No 863952 (ACE-OF-SPACE)). This work was also funded by the Max Planck Society, the EMBO Young Investigator Program and the Deutsche Forschungsgemeinschaft (DFG, German Research Foundation) under Germany’s Excellence Strategy – EXC 2117 – 422037984 and SFB1756 – 55955626. We are grateful to support from the Blue Sky research program of the University of Konstanz (Project EvoDevoGPT).

## Author contributions

Conceptualization: AL, LM, PM; Methodology: AL, GS, LL, SV, PM; Investigation: AL, LL; Visualization: AL, PM; Funding acquisition: PM; Project administration: PM; Writing: AL, PM. The authors declare that they have no competing interests.

## Materials and Methods

### Plasmid design and cloning

Plasmid sequences were designed using SnapGene or SeqBuilder. The sequences of the genetic parts were obtained from the iGEM parts repository (http://parts.igem.org/Catalog) and previous studies^32,34,65^. The plasmids were designed rationally based on the synthetic genetic circuit design. The circuit was implemented in parts on different plasmids to enable characterization of the output of each part. The AHL6 sensor plasmid pBC-A1-001 was a gift from Brian Chow^65^ (Addgene plasmid #78688, http://n2t.net/addgene:78688, RRID: Addgene_78688). The other plasmids were designed and either ordered as synthetic DNA fragments (Synbio Technologies) or cloned using standard molecular cloning techniques as described below.

The pAL101 plasmid was generated as follows. pTD103luxI_sfGFP^32^ (from Jeff Hasty, Addgene plasmid #48885, http://n2t.net/addgene:48885, RRID: Addgene_48885) was digested with EcoRI; the backbone containing *LuxI* and *LuxR* coding sequences was self-ligated and introduced into One Shot™ TOP10 chemically competent *E. coli* (Catalog number: C404010) by the heat-shock transformation method, and transformants were selected using kanamycin. The resulting plasmid was then modified by addition of a C-terminal 6xHis tag to *LuxI* using site-directed mutagenesis (Q5® Site-Directed Mutagenesis Kit - E0554S; primers 2985 and 2986, Supplementary Table 4). Correct clones were identified by Sanger sequencing (primers 2973, 2974, 2975 and 2976, Supplementary Table 4). Another site-directed mutagenesis was performed to insert a constitutive promoter (BBa_J23100, http://parts.igem.org/Part:BBa_J23100) upstream of *LuxR* (primers 2987 and 2988, Supplementary Table 4). The correct clones were identified by Sanger sequencing (primers 2973, 2974, 2975 and 2976, Supplementary Table 4). The plasmid was digested with ClaI and EcoRI and then assembled with DNA fragments containing AiiA (under the *pBAD* promoter) and sfGFP (under the *pLux* promoter) using the Gibson assembly method (Gibson Assembly® Cloning Kit - E5510S). The AiiA and sfGFP fragments were generated by PCR (AiiA fragment: [template: pTurL1, synthetic DNA (BioCat GmbH)] [primers 3086 and 3087, Supplementary Table 4], sfGFP fragment: [template: pTD103luxI_sfGFP –R, Addgene plasmid # 48887] [primers 3088 and 3089, Supplementary Table 4]). The resulting plasmid expresses sfGFP and LuxI under *pLux* promoters, LuxR under the constitutive promoter, and AiiA under the *pBAD* promoter. The plasmid sequence was confirmed using Sanger sequencing (primers 2973, 2976, 2977, 2978, 2496 and 2497, Supplementary Table 4).

The pAL102 plasmid was generated from the pAL101 plasmid by deletion of the *LuxR* cassette using the site-direction mutagenesis method (Q5® Site-Directed Mutagenesis Kit, primers 4093 and 4094, Supplementary Table 4).

The pAL103 plasmid was generated from the positive-feedback circuit (1) plasmid by restriction digestion (XmaI and EcoRI) and ligation with the insert containing a modified *pLux* promoter (primers 4466 and 4467, Supplementary Table 4). This resulted in a plasmid with a single stringent *pLux* promoter driving both *sfGFP* and *LuxI* bicistronically.

The pAL104 plasmid was generated from the pAL103 plasmid by deletion of the *LuxR* cassette using the site-direction mutagenesis method (Q5® Site-Directed Mutagenesis Kit, primers 4093 and 4094, Supplementary Table 4).

The entire sequence for the negative feedback circuit plasmids was ordered as two synthetic DNA fragments (from BioCat and Synbio Technologies) with compatible restriction sites (XhoI and AsiSI). The synthetic fragments were restriction-digested and ligated to generate the plasmid pAL201. Next, pAL201.1 was created from pAL201 by restriction digestion with PacI and self-ligation. pAL201.2 was created from pAL201.1 by modifying the ribosome-binding-site (RBS) upstream of *LasI* using site-directed-mutagenesis (primers 4201 and 4202, Supplementary Table 4). Spacer sequences flanking *LasR* cassette were added by PCR (primers 4254 and 4255, Supplementary Table 4), and the amplicon was cloned into the pAL201 backbone using FseI and XmaI sites to generate the plasmid pAL201.4.

The AHL12-sensor plasmid (pAL201.5) was created from pAL201.4. The coding sequence of *LasR* was PCR-amplified (primers 4275 and 4276, Supplementary Table 4) from pAL201.4 with flanking spacer^66^ sequences and restriction sites (AscI, XmaI); the pAL201.4 vector and the PCR amplified insert were restriction-digested with AscI and XmaI and ligated followed by transformation and selection of correct clones.

The AHL12 production plasmid (pAL209) was generated by performing multiple cloning steps as follows. The plasmid pAL205 was obtained by digesting pTD103aiiA(Cm)-R (a gift from Jeff Hasty, Addgene plasmid # 48888, http://n2t.net/addgene:48888, RRID:Addgene_48888) with XhoI and AvrII and ligated with *LasI-Myc*. The chloramphenicol resistance gene of pAL205 was replaced with the ampicillin resistance gene by restriction digestion and ligation to yield pAL206. The Myc-tag was deleted from LasI in pAL206 by site-directed mutagenesis (primers 4416 and 4417, Supplementary Table 4) to yield pAL209.

The LasR inhibition (1) plasmid (pAL210) and LasR inhibition (2) plasmid (pAL211) were generated as follows. A synthetic DNA fragment containing the coding sequence of anti-quorum-sensing protein 1 (Aqs1^53^) with flanking *pLux* promoter, RBS and terminator sequences was cloned into the pAL206 backbone using restriction digestion with XhoI and AvrII and ligation to obtain the pAL210 plasmid. The pAL210 plasmid was modified by site-directed mutagenesis (primers 4456 and 4457, Supplementary Table 4) to replace the *pLux* promoter and RBS, respectively, with a more stringent *pLux* and a weaker RBS, resulting in pAL211.

The AHL12 production + LasR inhibition plasmid (pAL214) was generated by combining pAL209 (AHL12 production, pTet:LasI) and pAL211 (LasR inhibition, *pLux*:Aqs1) plasmids. To this end, the pTet:LasI cassette was PCR-amplified (primers 4561 and 4562, Supplementary Table 4) from pAL209 and cloned into pAL211 using the XhoI restriction site.

The AHL12 sensor + AHL6 synthase plasmid (pAL215) was generated by inserting the AHL6 synthase gene under the *pLas* promoter (*pLas*:LuxI-FLAG from pAL201) into the pAL201.5 (AHL12 sensor) plasmid. To this end, the *pLas*:LuxI-FLAG cassette was PCR-amplified (primers 4554 and 4555, Supplementary Table 4) from pAL201 and cloned into pAL201.5 using the PacI restriction site. This plasmid contains a kanamycin resistance gene (*nptII*).

The plasmid pAL214_103 was generated by combining the positive feedback circuit parts form pAL103 with the pAL214 plasmid. To this end, the positive loop circuit sequences were PCR-amplified from pAL103 (primers 4586 and 4587, Supplementary Table 4) and cloned into pAL214 using the AatII restriction site. This resulted in the plasmid pAL214_103 with an ampicillin resistance marker and the circuit parts for positive feedback + AHL12 production + LasR inhibition.

The plasmid pAL214_104 was generated similar to the pAL214_103 plasmid. Here, pAL104 was used to PCR-amplify the positive loop parts. Note that pAL104 lacks the *LuxR* sequence.

### Generation of E. coli strains for characterization of the synthetic genetic circuits

*E. coli* (Marionette-sAJM.1504 strain from Christopher Voigt, Addgene #108251^54^) were transformed with the plasmid to be characterized using electroporation (voltage = 1.8 kV, 1 mm cuvette, BIO-RAD MicroPulser). The transformants were selected using an appropriate antibiotic. Glycerol stocks were prepared from overnight grown cultures (200 rpm, 37°C, ∼12 h) by mixing 0.5 ml of culture with 0.5 ml of 50% glycerol (v/v, aq.) and stored at -80°C.

Implementation of the synthetic circuits was achieved by transformation of *E. coli* with the plasmids listed in the table below:

| # | *Synthetic circuit strain* | *Plasmid(s)* | *Ori* | *Antibiotic* |
| --- | --- | --- | --- | --- |
| 1 | AHL<sub>6</sub> sensor (in DH5α strain) | pBC-A1-001<sup>65</sup> | pMB1 | Chloramphenicol |
| 2 | AHL<sub>12</sub> sensor | pAL201.5 | pMB1 | Kanamycin |
| 3 | Positive feedback circuit (1) | pAL101 | pMB1 | Kanamycin |
| 4 | Positive feedback circuit (2) | pAL102 | pMB1 | Kanamycin |
| 5 | Positive feedback circuit (3) | pAL103 | pMB1 | Kanamycin |
| 6 | Positive feedback circuit (4) | pAL104 | pMB1 | Kanamycin |
| 7 | AHL<sub>12</sub> production circuit | pAL209 | p15A | Ampicillin |
| 8 | LasR inhibition circuit (1) | pAL211 | p15A | Ampicillin |
| 9 | AHL<sub>12</sub> production and LasR inhibition (1) circuits | pAL214 | p15A | Ampicillin |
| 10 | AHL<sub>12</sub> sensor and AHL<sub>6</sub> production (via *pLas:LuxI-FLAG*) circuits | pAL215 | pMB1 | Kanamycin |
| 11 | AHL<sub>12</sub> production, LasR inhibition (1), and positive feedback (3) circuits | pAL214_103 | p15A | Ampicillin |
| 12 | AHL<sub>12</sub> production, LasR inhibition (1), and positive feedback (4) circuits | pAL214_104 | p15A | Ampicillin |
| 13 | AHL<sub>12</sub> sensor and AHL<sub>12</sub> production circuits | pAL201.5, pAL209 | pMB1, p15A | Kanamycin, Ampicillin |
| 14 | AHL<sub>12</sub> sensor and LasR inhibition (1) circuits | pAL201.5, pAL211 | pMB1, p15A | Kanamycin, Ampicillin |
| 15 | Negative loop [AHL<sub>12</sub> sensor, AHL<sub>12</sub> production and LasR inhibition (1)] circuit | pAL201.5, pAL214 | pMB1, p15A | Kanamycin, Ampicillin |
| 16 | Negative loop [AHL<sub>12</sub> sensor, AHL<sub>12</sub> production and LasR inhibition (1)] circuit + AHL<sub>6</sub> production (*pLas:LuxI-FLAG*) | pAL215, pAL214 | pMB1, p15A | Kanamycin, Ampicillin |
| 17 | Complete synthetic circuit (1)* | pAL214_103, pAL215 | p15A, pMB1 | Ampicillin, Kanamycin |
| 18 | Complete synthetic circuit (2) | pAL214_104, pAL215 | p15A, pMB1 | Ampicillin, Kanamycin |
| 19 | Constitutive mCherry circuit | pAL000 | pMB1 | Kanamycin |

\* The positive feedback module in the complete circuit is based on pAL103

### Fluorescence quantification in liquid culture

To analyze the dose-response behavior of engineered *E. coli* strains to treatment with different inducer doses in liquid cultures, fluorescence measurements were performed in 96-well plates. To this end, bacterial primary cultures were grown overnight (∼12 h) in LB medium at 37°C and 200 rpm (Sartorius Certomat® IS UHK-25), starting from single colonies or glycerol stocks. The next day, a secondary culture was inoculated and allowed to grow until early log phase (optical density at 600 nm (OD600) of 0.4). Cells were harvested by centrifugation (2000 rcf, Eppendorf centrifuge 5424) and re-suspended in fresh LB medium. The treatments were performed in 1.5 ml sterile tubes by adding appropriate amounts of inducer to 1 ml of the bacterial culture. The culture was transferred to 96-well plates (Corning® 3904). The fluorescence intensities and OD600 values were measured using a plate-reader (TECAN infinite M200 PRO).

### Diffusion coefficient measurements using sensor strains

Diffusion coefficients of the AHLs were determined in LB-agar at 37°C by adding a known amount of AHL to the lawn center of corresponding AHL sensor strains and analyzing the fluorescence gradient produced by the sensor strain. First, an overnight primary culture and a subsequent secondary culture of the appropriate AHL sensor strain was grown as described above. Then, cells were harvested by centrifugation and re-suspended in LB medium to an effective OD600 of 1.0. This resuspension was used to coat a plate with a ∼1 mm layer of LB-agar and grown at 37°C for 4 h to generate a near-uniform bacterial lawn. A known amount of corresponding AHL (1 µl of 1 µM) was added to the center of the lawn to induce expression of a fluorescent reporter (sfGFP for AHL6 and mCherry for AHL12). Images were acquired every hour (0 h to 4 h) using an Axio Zoom V16 microscope (ZEISS). Control measurements with fluorescent dextrans (3 kDa and 10 kDa) were performed using the same setup.

Radial profiles of fluorescence intensity were measured using Fiji^67^ (Radial Profile plugin) and saved as .csv files. A custom Python script was developed (https://github.com/mueller-lab/SyntheticPatterns/) to generate simulated datasets and determine the diffusion coefficients by fitting the simulated profiles to the experimental data. The noisy ends of the gradients were omitted while generating the fits. The mathematical models were based on the experimental data of the reaction kinetics in liquid cultures obtained using plate-reader assays. FiPy^68^ was used to solve partial differential equations (PDEs) describing the mathematical models. The SciPy^69^ library of Python (scipy.optimize.minimize) was used for solving the optimization problem to get the best fit. This method may slightly underestimate the diffusion rates because the geometry of the system in the numerical simulations is one-dimensional, whereas in the experimental setup the molecules diffuse in a 3D agar slab (diameter ∼35 mm, thickness ∼1 mm). Note that the fluorescent reporters used in AHL6 sensor and AHL12 sensor strains have different maturation kinetics.

### Fluorescence imaging of bacterial lawn

A quasi-2D time-lapse imaging method was developed for the analysis of spatiotemporal patterning in a multicellular lawn of synthetic bacterial strains. First, the bacteria were grown overnight in LB medium containing the proper antibiotic at 200 rpm and 37°C. The next day, a secondary culture was started by inoculating the overnight culture in plain LB medium (1/100 dilution). After the OD600 of 0.4 was reached, the cells were harvested by centrifugation and re-suspended in a gel of 0.5% low-melting-temperature agarose (NuSieve™ GTG™ Agarose, Catalog #: 50080) to make a cell-agarose mixture with 5-fold higher cell density than that of the 0.4 OD600 culture. This cell mixture was then loaded carefully on LB-agar in a 24-well plate to get a lawn of bacteria. The mild curvature of the agar surface causes a slightly higher initial cell-density in the middle of the well (Supplementary Fig. 10a-d). The cells were allowed to grow for two hours before making treatments and starting the time-lapse imaging.

Time-lapse imaging was performed using an automated high-throughput fluorescence microscope (Keyence BZ-X810) at 15 min intervals. Images were acquired using a 2x apochromat objective (PlanApo 2x 0.10/8.50mm) with a 3.7 W LED transmitted light source, and an 80 W metal halide lamp or 40 W LED as fluorescent light source. Images were saved as 8-bit, 480 pixel × 360 pixel or 1920 pixel × 1440 pixel TIFF files at a resolution of 15 µm/pixel.

### Image analysis

Image analysis was performed using Fiji^67^ (ImageJ 2.9.0/1.53t). Image stitching and quantifications were performed using a custom ImageJ macro (https://github.com/mueller-lab/SyntheticPatterns). The quantifications of mean intensity and radial fluorescence profiles were saved as .csv files and plotted using custom Python code (https://github.com/mueller-lab/SyntheticPatterns). The BaSiC ImageJ plugin was used for shading correction^70^. Gray value ranges of the 16-bit images were adjusted for better visualization (Fig. 3: sfGFP 10-255, mCherry 20-60). *viridis* and *magma* lookup tables were applied to sfGFP and mCherry images, respectively.

### Quantitative reverse transcription PCR (qRT-PCR)

Bacterial cells were harvested as described in the *Fluorescence quantification in liquid culture* section. The treatments were performed in 1.5 ml sterile tubes by adding appropriate amounts of inducer to 1 ml of the bacterial culture, which was incubated for 4 h at 37°C and 200 rpm. The cells were pelleted at 2000 rcf with a standard tabletop centrifuge (Eppendorf 5424). Total RNA was extracted using NucleoZOL (#740404.200). Synthesis of cDNA was performed using up to 1 µg of total RNA and the SuperScript™ III First-Strand Synthesis SuperMix (#11752050) kit. No-reverse transcription control reactions were performed in parallel. The cDNA was 5-fold diluted with nuclease-free water and used to set up the qPCR reaction with appropriate primers (Supplementary Table 4) and the Platinum™ SYBR™ Green qPCR SuperMix-UDG kit (#11733038). The qPCR reaction and quantification were performed using the CFX Connect Real-Time PCR Detection System. Data analysis and plotting was performed using Python (https://github.com/mueller-lab/SyntheticPatterns). The efficiencies of all primers were validated in dilution experiments (Supplementary Fig. 7).

### Mathematical modeling

Mathematical models were developed to numerically simulate system dynamics using known interactions and reaction kinetics and also to predict the system behavior upon perturbations. All equations are described in the Supplementary Note, and parameter values are provided in Supplementary Table 1. All numerical simulations were performed by implementing the finite volume PDE solver FiPy^68^ in custom Python code (available at https://github.com/mueller-lab/SyntheticPatterns). The simulations were performed on a discrete 1D or 2D grid with zero-flux boundary conditions. The simulation results for Fig. 3, Fig. 4 and Supplementary Fig. 10-11 were generated by performing 1D simulations and setting the simulation conditions (initial and boundary conditions) as closely matching the experimental conditions as possible. These findings were further confirmed in 2D simulations.

## Data and code availability

The open-source code along with example data is available from https://github.com/mueller-lab/SyntheticPatterns.

