## Supplementary Information for "Self-organized traveling waves in a synthetic multicellular reaction-diffusion system"

**Contents**

Supplementary Note

Supplementary Tables 1-4

Supplementary Figures 1-11

Legends for Supplementary Movies 1-8

Supplementary References

### Supplementary Note

#### *Mathematical modeling*

Following the initial design of the patterning circuit and coarse-grained mathematical simulations, we built refined mathematical models in line with our experimental approach to design and implement the modular circuits.

First, we consider the AHL sensor module:

*Model 1: $AHL_6$ sensor*

The following system of partial differential equations (PDEs) describes the $AHL_6$ sensor model. The sensor works by activation of the *pLux* promoter through cooperative binding of $AHL_6$ and LuxR. The cooperativity is implemented by using a Hill-type function for LuxR-$AHL_6$ complex formation, $H(X, Y, K_T, nH, k_{leak})$:

$$H(X, Y, K_T, nH, k_{leak}) = X\left(k_{leak} + \frac{Y^{nH}}{K_T^{nH} + Y^{nH}}\right); 1 \leq nH \leq 3;$$

$$\frac{\partial}{\partial t}LuxR = k_{prot} - k_{d_{prot}}LuxR$$

$$\frac{\partial}{\partial t}AHL_6 = -k_{d_{ahl}}AHL_6 + D_6\nabla^2 AHL_6$$

$$\frac{\partial}{\partial t}GFP = k_{gfp}LuxR_{AHL6} - k_{d_{ssrA}}GFP$$

with $LuxR_{AHL6} = H(LuxR, AHL_6, K_{T6}, nH_6, k_{leak})$. $AHL_6$ was varied in the range [0,1000] nM.

Parameter values for this model were determined from our experimental data (Supplementary Fig. 8a-d). For some parameters, reported estimates from the literature were used[1] (Supplementary Table 1). The model was simulated using the FiPy solver[2], and the results are shown in Supplementary Fig. 8e,f.

*Model 2: $AHL_{12}$ sensor*

This model describes the dose-response behavior of the $AHL_{12}$ sensor, similar to Model 1:

$$\frac{\partial}{\partial t}LasR = k_{prot} - k_{d_{prot}}LasR$$

$$\frac{\partial}{\partial t}AHL_{12} = -k_{d_{ahl}}AHL_{12} + D_{12}\nabla^2 AHL_{12}$$

$$\frac{\partial}{\partial t}mCherry = k_{mChr}LasR_{AHL12} - k_{d_{ssrA}}mCherry$$

with $LasR_{AHL12} = H(LasR, AHL_{12}, K_{T12}, nH_{12}, k_{leak})$. $AHL_{12}$ was varied in the range [0.01,1000] nM.

The results of numerical simulations of this mathematical model are shown in Supplementary Fig. 6e.

*Model 3: Positive feedback module*

This mathematical model describes the positive feedback loop implemented by expressing the gene responsible for $AHL_6$ production, i.e. *LuxI*, under the $AHL_6$-inducible *pLux* promoter. The model is an extension of Model 1 that includes *pLux*-driven $AHL_6$ production and arabinose-driven $AHL_6$ degradation:

$$\frac{\partial}{\partial t}LuxR = k_{prot} - k_{d_{prot}}LuxR$$

$$\frac{\partial}{\partial t}AHL_6 = k_{ahl}LuxI - AHL_6\left(k_{d_{ahl}} + k_{d1_{ahl}}AiiA\right) + D_6\nabla^2 AHL_6$$

$$\frac{\partial}{\partial t}sfGFP = k_{sfGFP}LuxR_{AHL6} - k_{d_{ssrA}}sfGFP$$

$$\frac{\partial}{\partial t}AiiA = k_{prot}ara_{indc} - k_{d_{ssrA}}AiiA$$

$$\frac{\partial}{\partial t}LuxI = k_{prot}LuxR_{AHL6} - k_{d_{ssrA}}LuxI$$

with $LuxR_{AHL6} = H(LuxR, AHL_6, K_{T6}, nH_6, k_{leak})$. This Hill-type term controls production of LuxI and sfGFP in $AHL_6$ concentration-dependent manner.

$ara_{indc} = H(10, ara, K_{Tara}, nH_{ara}, k_{leak})$. *ara* was varied in the range [0, $10^6$].

The simulation results are shown in Fig. 2d. The parameter values are provided in Supplementary Table 1.

*Model 4: Negative feedback module ($AHL_{12}$ sensor + LasR inhibition)*

The negative feedback was implemented by expression of the LasR inhibitor Aqs1 under the *pLux* promoter. This model simulates the behavior of the $AHL_{12}$ sensor and the LasR inhibition circuit:

$$\frac{\partial}{\partial t}LasR = k_{prot} - k_{d_{prot}}LasR$$

$$\frac{\partial}{\partial t}mCherry = k_{mCherry}LasR_{AHL12} - k_{d_{ssrA}}mCherry$$

$$\frac{\partial}{\partial t}Aqs1 = k_{prot}(LuxR_{AHL6} + LuxR_{AHL12}) - k_{d_{prot}}Aqs1$$

with $LasR_{AHL12} = H(LasR, AHL_{12}, K^*_{T12}, nH_{12}, k_{leak})$, where $K^*_{T12}$ is given by

$$K^*_{T12} = \frac{K_{T12}}{H\left(1.0, K_{Taqs}, Aqs1, nH_{aqs}, k_{leak}\right)}$$

implementing the Aqs1-mediated inhibition of LasR-$AHL_{12}$. $AHL_{12}$ levels were varied in the range [0.1,1000] similar to the dose-response assay.

$LuxR_{AHL6} = H(LuxR, AHL_6,\ K_{T6}, nH_6,\ k_{leak})$; AHL6 was set to 0 nM or 1000 nM.

$LuxR = 10$ is assumed to be at the steady-state level. *LuxR* was expressed from the Marionette strain chromosome from a constitutive promoter.

$LuxR_{AHL12} = H(LuxR, AHL_{12}, K_{T12xR}, nH_{12}, k_{leak})$ gives the non-specific activation of the *pLux* promoter by binding of LuxR and AHL12. The simulation results are shown in Supplementary Fig. 6k. The parameter values are provided in Supplementary Table 1.

Next, we consider the spatiotemporal dynamics of the complete synthetic multicellular system by combining the positive and negative feedback modules.

*Model 5: Complete synthetic circuit*

To explain the emergence of traveling wave patterns in the experimental system and make predictions about how the patterns can be modified with different perturbations, we developed a mathematical model of the complete synthetic circuit. The PDE-based mathematical model considers key variables and parameters of the system to capture important system properties. Model 3 and Model 4 were used as a basis for modeling the feedback interactions. The diffusion coefficients for AHLs were set to experimentally determined values. The variables *LuxR*, *LasR*, and *LasI* were set to constant values to reduce model complexity without losing important dynamical aspects. The role of cell growth and nutrient depletion was considered using the variable $cell_D$.

$$\frac{\partial}{\partial t} LuxI = cell_D\left(k_{prot}(pLux + pLas) - k_{d_{ssrA}} LuxI\right)$$

$$\frac{\partial}{\partial t} AHL_{12} = cell_D\left(k_{ahl} AHL_{12cap} LasI - k_{d_{ahl12}} AHL_{12} AiiA\right) - k_{d_{AHL}} AHL_{12} + D_{12} \nabla^2 AHL_{12}$$

$$\frac{\partial}{\partial t} AHL_6 = cell_D\left(k_{ahl} AHL_{6cap} LuxI - k_{d_{ahl6}} AHL_6 AiiA\right) - k_{d_{AHL}} AHL_6 + D_6 \nabla^2 AHL_6$$

$$\frac{\partial}{\partial t} mCherry = cell_D\left(k_{mChr} pLas - k_{d_{ssrA}} mCherry\right)$$

$$\frac{\partial}{\partial t} sfGFP = cell_D\left(k_{gfp} pLux - k_{d_{ssrA}} sfGFP\right)$$

$$\frac{\partial}{\partial t} AiiA = cell_D\left(k_{prot} ara_{indc} - k_{d_{ssrA}} AiiA\right)$$

$$\frac{\partial}{\partial t} Aqs1 = cell_D\left(k_{prot} pLux - k_{d_{prot}} Aqs1\right)$$

$$\frac{\partial}{\partial t} cell_D = cell_D\left(\frac{k_{cell}}{K_{cell}}\right)(K_{cell} - cell_D) + D_c \nabla^2\ cell_D$$

with $LasR_{AHL12} = H(LasR, AHL_{12}, K^*_{T12}, nH_{12}, k_{leak})$, where $K^*_{T12}$ is given by

$K^{*}_{T12} = \frac{K_{T12}}{H(1.0, K_{Taqs}, Aqs1, nH_{aqs}, k_{leak})}$ as in Model 4

$LuxR_{AHL6} = H(LuxR, AHL_6, K_{T6}, k_{leak})$ as in Model 3

$LuxR_{AHL12} = H(LuxR, AHL_{12}, K_{T12xR}, k_{leak})$

$LuxR = 40, LasR = 40, LasI = 2000$

$pLux = LuxR_{AHL6} + LuxR_{AHL12}$

$pLas = LasR_{AHL12}$

$ara_{indc} = H(10, ara, K_{Tara}, nH_{ara}, k_{leak})$ as in Model 3

$ara \in (0, 1, 10^3, 10^6)$

To implement limits on AHL production due to nutrient depletion after cell growth, the following terms were applied:

$AHL_{6cap} = \frac{(K_{cell} - cell_D)}{lawnFrac * K_{cell}}$;

$AHL_{12cap} = \frac{(1.1 * K_{cell} - cell_D)}{1.1 * lawnFrac * K_{cell}}$; the factor 1.1 is used to allow some $AHL_{12}$ production at higher cell density (Supplementary Fig. 10a-d).

The parameter $lawnFrac$ (between 0 to 1) controls the fraction of the domain occupied by cells. In 1D simulations it equals the length fraction of domain occupied by cells, and in 2D simulations it equals the area fraction of the domain occupied by cells.

Note that not all parameters could be precisely measured and were therefore set to biologically reasonable values capable of recapitulating our experimental observations (Supplementary Table 1).

***Prospects and potential for future work***

Here, we outline experiments and extensions that could build on our synthetic multicellular patterning system. A main limitation of the current experimental setup is that the dynamics reach a steady state after approximately 24 h, after which no further wave propagation is observed. Our data and model suggest that this behavior likely reflects a combination of nutrient limitation, reduced cellular resources, metabolic burden imposed by the multicomponent circuit, altered AHL production in dense cultures, and dilution or loss of AHL signals into the surrounding agar. Limiting levels of S-adenosyl methionine (SAM), a precursor for AHL synthesis also important for cell division[3], may contribute to these effects since nutrient depletion due to cell division and growth may lead to reduced rates of AHL production. However, the loss of sfGFP signal in the positive-feedback-only system suggests that SAM competition between LuxI and LasI is unlikely to be the sole explanation. Future experiments using richer nutrient conditions, supplementation with SAM, or strains optimized for reduced metabolic interference could therefore help determine which physiological constraints limit long-term circuit dynamics.

Another important direction will be to analyze the system at higher spatial and cellular resolution. In the present study, live imaging was performed at the scale of entire bacterial lawns, which enabled quantification of multicellular spatiotemporal dynamics but did not resolve single-cell gene-expression variability or local growth dynamics. Single-cell-resolution imaging would make it possible to determine how stochastic gene-expression events, local differences in cell density, and cell division within the agarose contribute to the emergence and propagation of lawn-scale patterns. Such experiments could also be combined with stochastic modeling to complement the deterministic PDE model used here.

Further work should also directly quantify spatial features of the initial bacterial lawn. In the current setup, the shallow cell-density profile is inferred from the preparation procedure, bright-field image analysis, and the ability of the model to recapitulate the observed dynamics. Higher-resolution bright-field profiling, independent density calibration, or CFU-based sampling would provide a more direct estimate of the initial cell-density distribution and help refine the model assumptions.

There is also substantial potential to expand the accessible parameter space of the circuit. In the negative-feedback branch, basal pTet activity is sufficient to drive LasI expression and $AHL_{12}$ production, while anhydrotetracycline induction provides only a limited dynamic range in mCherry output. Optimizing this part of the circuit, for example by using alternative promoters, receptor variants, degradation tags, or chromosomal integration strategies, could improve tunability and may allow access to additional dynamical regimes. A more systematic dissection of circuit variants, including removal or rewiring of individual positive- and negative-feedback branches, would further clarify how each module contributes to amplification, timing, regularity, and wave propagation.

Future versions of the system could also include spatially controlled inputs. For example, optogenetic activation or local inducer application would make it possible to initiate signaling from defined positions and thereby separate induced wave propagation from spontaneous pattern emergence caused by cell-density heterogeneity or local fluctuations. This would be particularly useful for testing whether and under which conditions the system can support externally triggered propagation fronts.

Finally, the circuit could be improved by minimizing unintended interactions with host-cell physiology and endogenous or strain-encoded AHL-responsive components. Although the current module-level characterization supports the intended function of the implemented circuit, future work in alternative *E. coli* backgrounds, including strains with reduced AHL cross-talk and optimized transcriptional regulators, would further strengthen the orthogonality of the system. Such optimized implementations may enable more robust long-term patterning, more precise quantitative control, and the engineering of additional multicellular outputs beyond fluorescent reporters[4,5].

**Supplementary Table 1. Parameters used in the simulations.**

| Parameter | Symbol | Value | Reference |
|---|---|---|---|
| Protein synthesis rate | $k_{prot}$ | 0.2 $min^{-1}$ | |
| sfGFP synthesis and maturation rate | $k_{sfGFP}$ | 0.1 $min^{-1}$ | |
| mCherry synthesis and maturation rate | $k_{mCherry}$ | 0.02 $min^{-1}$ | |
| Protein degradation rate | $k_{dprot}$ | 0.0028 $min^{-1}$ | |
| LVA-tagged protein degradation rate | $k_{dssrA}$ | 0.018 $min^{-1}$ | [6] |
| AHL synthesis rate (via LuxI) | $k_{ahl}$ | 0.002 $min^{-1}$ | |
| AHL decay rate (hydrolysis) | $k_{dahl}$ | 0.0011 $min^{-1}$ (Half-life ~10 h) | [1,7] |
| $AHL_6$ degradation rate (by Lactonase) | $k_{dahl6}$ | 0.01 $min^{-1}$ | [8] |
| $AHL_{12}$ degradation rate (by Lactonase) | $k_{dahl12}$ | 0.001 $min^{-1}$ | [8] |
| $AHL_6$ activation threshold | $K_{T6}$ | 10 nM | This study |
| Hill coefficient for $AHL_6$ | $nH_6$ | 1.5 | This study |
| $AHL_6$ diffusion coefficient | $D_6$ | 350 $\mu m^2/s$ | This study |
| $AHL_{12}$ activation threshold | $K_{T12}$ | [50, 80] nM | This study |
| $AHL_{12}$ threshold concentration for LuxR binding | $K_{T12xR}$ | [200, 2000] nM | |
| Hill coefficient for $AHL_{12}$ | $nH_{12}$ | 2 | This study |
| $AHL_{12}$ diffusion coefficient | $D_{12}$ | 350 $\mu m^2/s$ | This study |
| Arabinose threshold concentration | $K_{Tara}$ | 6000 nM | This study |
| Hill coefficient for arabinose | $nH_{ara}$ | 1.5 | This study |
| Aqs1 threshold concentration | $K_{Taqs}$ | [200, 300] nM | |
| Hill coefficient for Aqs1 | $nH_{aqs}$ | 2 | |
| Promoter leakiness | $k_{leak}$ | [0.05, 0.08] | |
| Cell density increase rate | $k_{cell}$ | 0.007 | |

| | | |
|---|---|---|
| Cell density capacity | $K_{cell}$ | 1.6 |
| Cell movement (diffusion) constant | $D_c$ | 0.1 μm$^2$/s |
| Lawn fraction, i.e. the fraction of the domain covered with cells | *lawnFrac* | 0.5, 0.65, 1 (for 1D) |

**Supplementary Table 2. Arabinose threshold concentrations for the positive feedback module.**

| Strain | Arabinose threshold (nM) | Treatment time (min) | Dynamic range |
|---|---|---|---|
| pAL101 | 967 ± 40 | 360 | 4.2 |
| pAL102 | 1402 ± 41 | 120 | 2.4 |
| pAL103 | 480 ± 18 | 360 | 6.5 |
| pAL104 | 880 ± 37 | 120 | 3.9 |

**Supplementary Table 3. Diffusion coefficient measurements.**

| Molecule | ***D*** **[mean ± stdev] (µm²/s)** | **Samples (*n*)** | **Previous estimates (µm²/s)** | **Reference** |
|---|---|---|---|---|
| Dextran (3 kDa) | 120 ± 5 | 7 | 170 ± 22 | [9] |
| Dextran (10 kDa) | 67 ± 12 | 7 | 83 ± 8 | [9] |
| $AHL_6$ | 313 ± 69 | 12 | 17 | [10] |
| $AHL_{12}$ | 320 ± 59 | 11 | 83 | [11] |

Note: Previous estimates listed here are from different experimental setups and conditions.

**Supplementary Table 4. Primers used in this study.**

| Name | Sequence |
|---|---|
| 2985_His_SDM_F | atcaccatgccgcagcagcgaacgacgaaaat |
| 2986_His_SDM_R | ggtgatggtgtgaatttaagactgctttttaaactgttc |
| 2973_LuxR_qPCR_F | catacggctaacaatggcttcg |
| 2974_LuxR_qPCR_R | gcatgcccacgctaaacatt |
| 2975_LuxI-ssrA_qPCR_F | ggtcaacagagtgctcccaa |
| 2976_LuxI-ssrA_qPCR_R | tcgctctattgctgttgatgttac |
| 2987_pJ23_SDM_F | agctagccgtcaaatcgataattgtcccgggaattgtgatatcacctattgtttgtcgcaag |
| 2988_pJ23_SDM_R | cagtcctaggtacagtgctagcggatccaaagaggagaaacatatgaaaaacataaatgccgac |
| 3086_Gibson_aiiA_F | gactgagctagccgtcaaatcgataggttctgttaagtaactgaacccaatgtcgttagtgacgcttacctcttaagaggtcactgacctaacaacattgattatttgcacggc |
| 3087_Gibson_aiiA_R | tataacaaaccattttcttgcgtaaacctgtacgatcctacaggtacaattcccggggagagcgttcaccgac |
| 3088_Gibson_sfGFP_F | caagaaaatggtttgttatagtcgaataaagcgatcgcattaaagaggagaaaggtaccatgag |
| 3089_Gibson_sfGFP_R | tttctcctctttaatgaattctttattcgactataacaaaccattttcttgcgtaaacctgtacgatcctacaggtgatatcatttgtcctactcaggagagc |
| 2977_sfGFP_qPCR_F | tacaagacgcgtgctgaagt |
| 2978_sfGFP_qPCR_R | ccatcttcaacgttgtggcg |
| 2979_aiiA-LAA_qPCR_F | agcgaacggaatatgaggca |
| 2980_aiiA-LAA_qPCR_R | cgactgatggcctggagaat |
| 2496_aiiA_F | aaagagtgcatcctgccgaa |
| 2497_aiiA_R | ggggtatgtccaggcgtatg |
| 4093_SDM_Mar_F | atcgataggttctgttaagtaac |
| 4094_SDM_Mar_R | gtgaagacgaaagggc |
| 4466_pAL103Lux_F | cgctctccccgggaattgtacctgtaggatcgtacaggtttacgcaagaaaatggtttgttacagtcgaataaagcgatcgc |
| 4467_pAL103Lux_R | ctctttaatgaattcttacgctgcaagggcg |
| 4201_SDM_201_pTet_F | ggggaaatactagatgattgttcagattggc |
| 4202_SDM_201_pTet_R | tctttctctagtattaaacaaaattatttgtagaggc |

| | |
|---|---|
| 4254_pAL201_4_F | atgatcggccggccgacattgggttcagttacttaacagaacctttgacggctagctcagtcctaggtacagtgctagcggatccaaagaggagaaacatatggccttggttgacgg |
| 4255_pAL201_4_R | atgatccccgggaggttctgttaagtaactgaacccaatgtcgttagtgacgcttacctcttaagaggtcactgacctaacagagagcgttcaccgacaaac |
| 4275_pAL201_5_F | atgatcggcgcgcccctgcagtattcattttcagcttacggaaggtagatttgacggctagctcagtc |
| 4276_pAL201_5_R | atgatccccgggcgctgcgtctgcctcctgaccacttgagtaactgctgcgaaaaaaccccgccgaagcggggtttttttgcgtcaacgcgtcgcaaac |
| 4352_LasI_pAL205_F | agatgatcctcgagagatgctgtagtgggatg |
| 4353_LasI_pAL205_R | agatgatccctaggttaaacaaaattatttgtagaggactg |
| 4416_LasI_SDM_F | taacgccgaaaaccccgc |
| 4417_LasI_SDM_R | ggaaacggccaggcgttg |
| 4456_pAL211_SDM_F | aaagacatgagttactagatgaccaacaccgatctg |
| 4457_pAL211_SDM_R | ctctagatttattcgactgtaacaaaccattttcttgcg |
| 4458_pAL212_SDM_F | aaagacatgagttactagatgagtaaaaaaatcaaactgc |
| 4459_pAL212_SDM_R | ctctagatttattcgactgtaacaaaccattttcttgc |
| 4561_pAL214_x_lasI_F | agatgatcctcgagagatgctgtagtg |
| 4562_pAL214_x_lasI_R | agatgatcctcgagttaaacaaaattatttgtagaggactg |
| 4554_pAL215_luxI_F | agatgatcttaattaagagaaagaggagaaatactagatgacg |
| 4555_pAL215_luxI_R | agatgatcttaattaattacttgtcgtcatcgtctttg |
| 4586_posLoop_F | agatgatcgacgtcaaataggcgtatcacgagg |
| 4587_posLoop_R | agatgatcgacgtcaccgtattaccgcctttg |

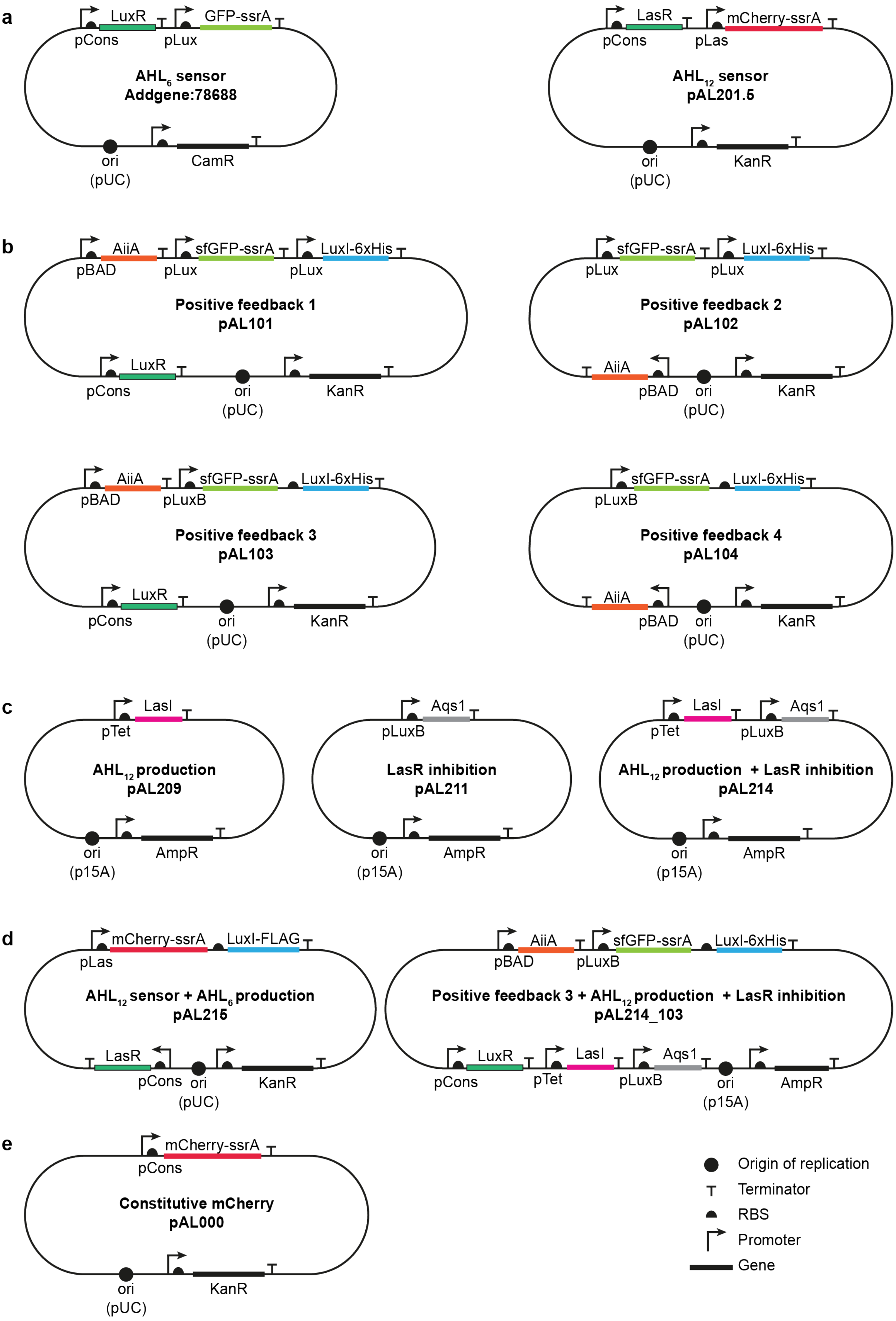

**Supplementary Figure 1 | Plasmid schematics. (a)** AHL sensor plasmids. **(b)** Positive feedback module plasmids: pAL101 – constitutive *LuxR* and less stringent *pLux*, pAL102 –

*LuxR* removed and less stringent *pLux*, pAL103 – constitutive *LuxR* and more stringent *pLux*, pAL104 – *LuxR* removed and more stringent *pLux*. **(c)** Negative feedback module plasmids: pAL209 – $AHL_{12}$ production, pAL211 – LasR inhibition, pAL214 – $AHL_{12}$ production and LasR inhibition. **(d)** Complete synthetic circuit plasmids: pAL215 – $AHL_{12}$ sensor and $AHL_6$ production, pAL214_103: pAL103 cloned into pAL214. **(e)** Constitutive mCherry expression plasmid.

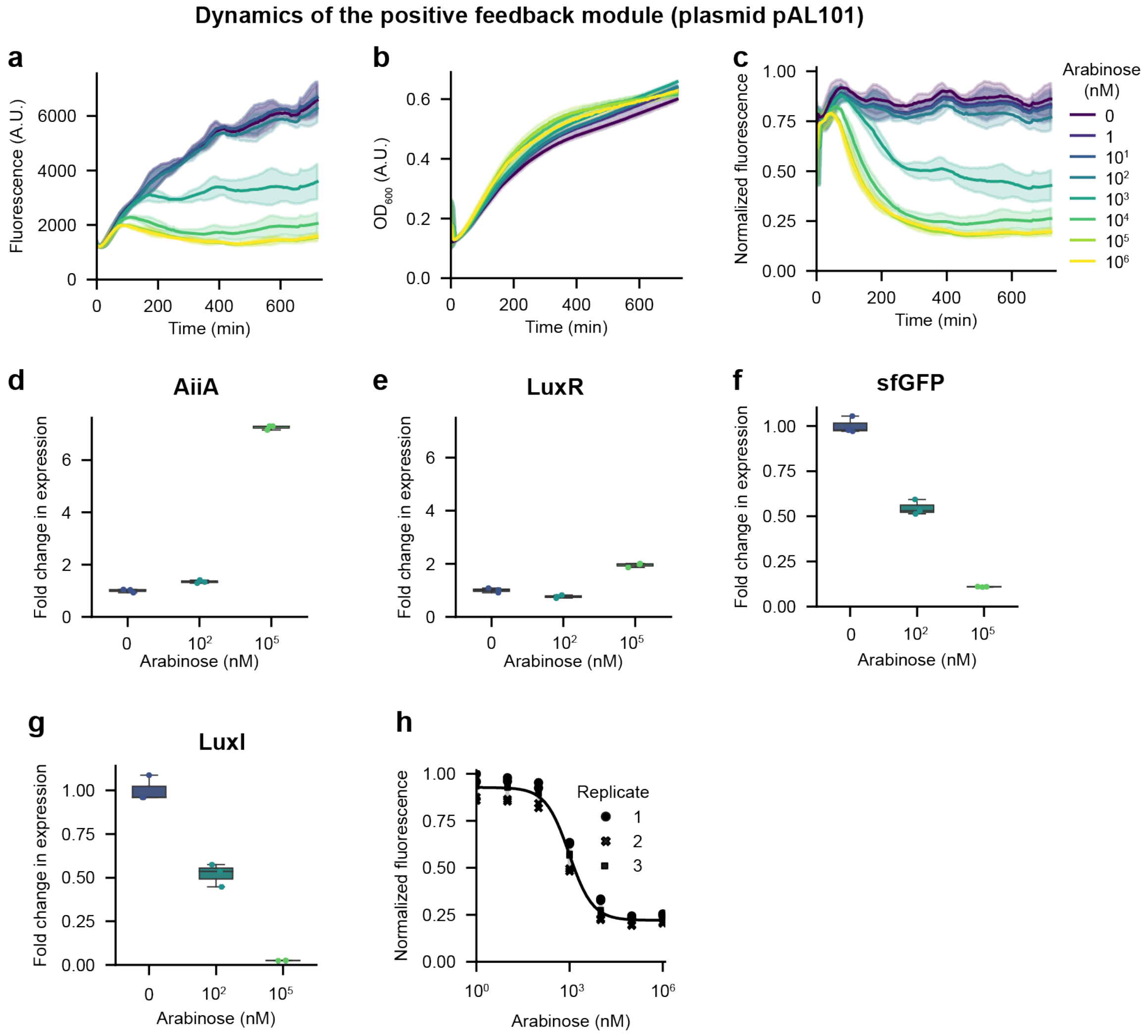

**Supplementary Figure 2 | Characterization of the positive feedback module (plasmid pAL101). (a-c)** Time-series plots showing sfGFP fluorescence (a), $OD_{600}$ (b), and normalized fluorescence/$OD_{600}$ (c) for the positive feedback strain carrying plasmid pAL101 with different arabinose treatments. n = 3 biological replicates with 2 technical replicates each. Error envelopes indicate standard deviation. **(d-g)** Boxplots showing fold-change in mRNA levels of *AiiA*, *LuxR*, *sfGFP*, and *LuxI* quantified by qRT-PCR after 4 h arabinose treatments. n = 1 biological replicate with 3 technical replicates. The center line represents the median, while the upper and lower box edges correspond to the first and third quartiles, respectively. Whiskers extend to the farthest data point within 1.5 times the interquartile range from the box edges or to the maximum data point within this range, whichever is closer. *ihfβ* was used as a reference gene for all RT-qPCR analyses. **(h)** Dose-response curve at 6 h after treatment with arabinose for the data shown in (a-c). Error bars indicate standard deviation.

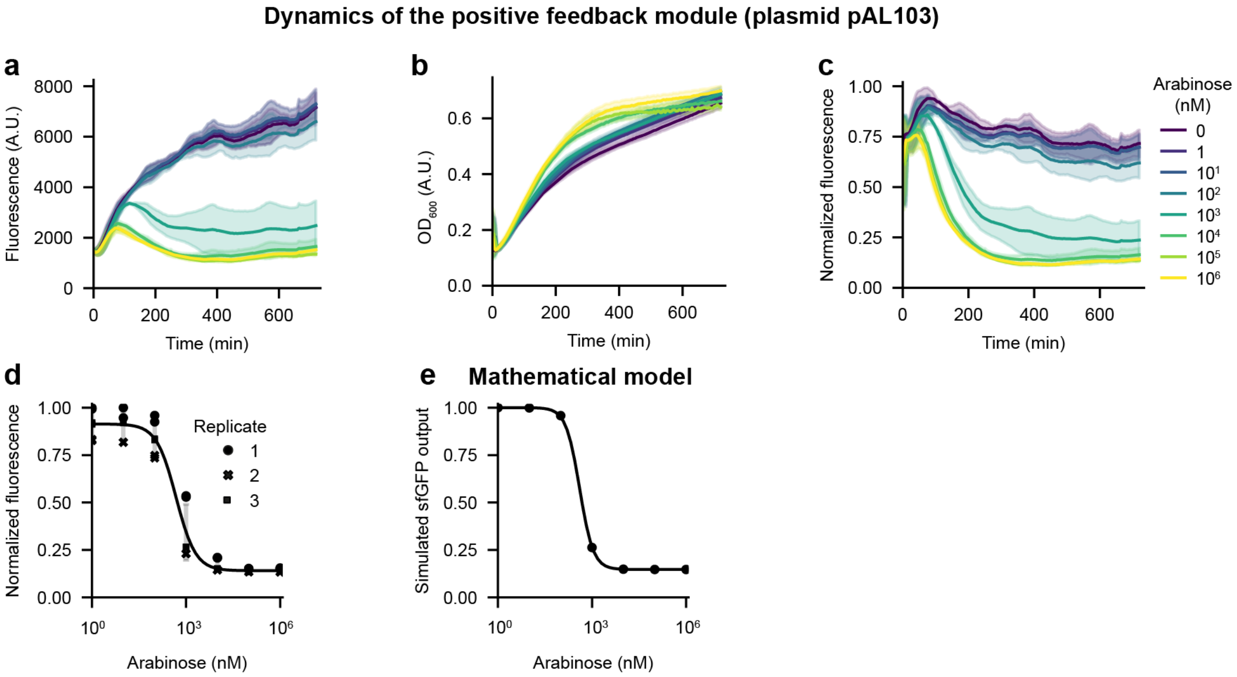

**Supplementary Figure 3 | Characterization of the positive feedback module (plasmid pAL103). (a-c)** Time-series plots showing sfGFP fluorescence (a), $OD_{600}$ (b), and normalized fluorescence/$OD_{600}$ (c) for the positive feedback strain carrying plasmid pAL103 with different arabinose treatments. n = 3 biological replicates with 2 technical replicates each. Error envelopes indicate standard deviation. **(d)** Dose-response curve at 6 h after treatment with arabinose for the data shown in (a-c). A reverse-sigmoid function was fitted to the data to determine the arabinose threshold concentration (480 ± 18 nM). Error bars indicate standard deviation. **(e)** A simulated dose-response curve using the mathematical model for the positive feedback module. Data in (c) is the same as in Fig. 2c.

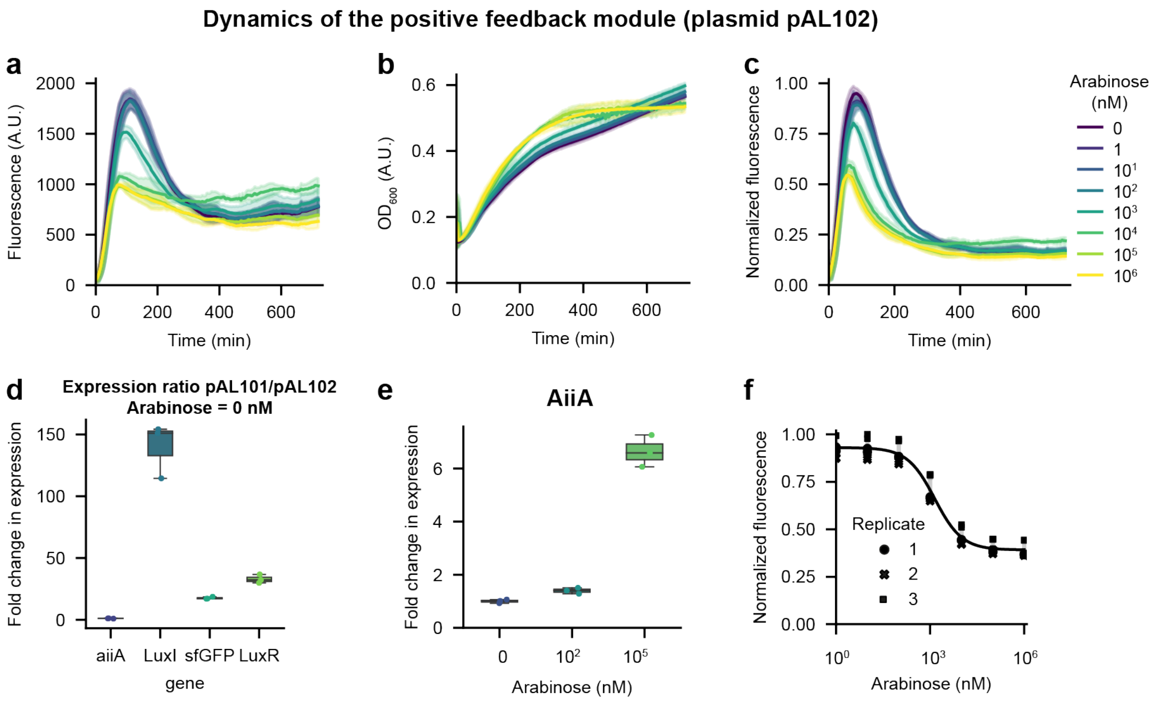

**Supplementary Figure 4 | Characterization of the positive feedback module (plasmid pAL102). (a-c)** Time-series plots showing sfGFP fluorescence (a), $OD_{600}$ (b), and normalized fluorescence/$OD_{600}$ (c) for the positive feedback strain carrying plasmid pAL102 with different arabinose treatments. n = 3 biological replicates with 2 technical replicates each. Error envelopes indicate standard deviation. **(d)** Relative expression of *AiiA*, *LuxI*, *sfGFP* and *LuxR* in pAL101 and pAL102 strains without arabinose treatments. The strain carrying pAL102 only has a single chromosomal copy of *LuxR*, while the pAL101 strain also expresses *LuxR* from the plasmid. Reference gene *ihfβ* for all qRT-PCR analyses. **(e)** Boxplots showing fold-change in mRNA levels of *AiiA* quantified by qRT-PCR after 4 h arabinose treatments. n = 1 biological replicate with 3 technical replicates. The center line represents the median, while the upper and lower box edges correspond to the first and third quartiles, respectively. Whiskers extend to the farthest data point within 1.5 times the interquartile range from the box edges or to the maximum data point within this range, whichever is closer. **(f)** Dose-response curve at 3 h after treatment with arabinose for the data shown in (a-c). Error bars indicate standard deviation.

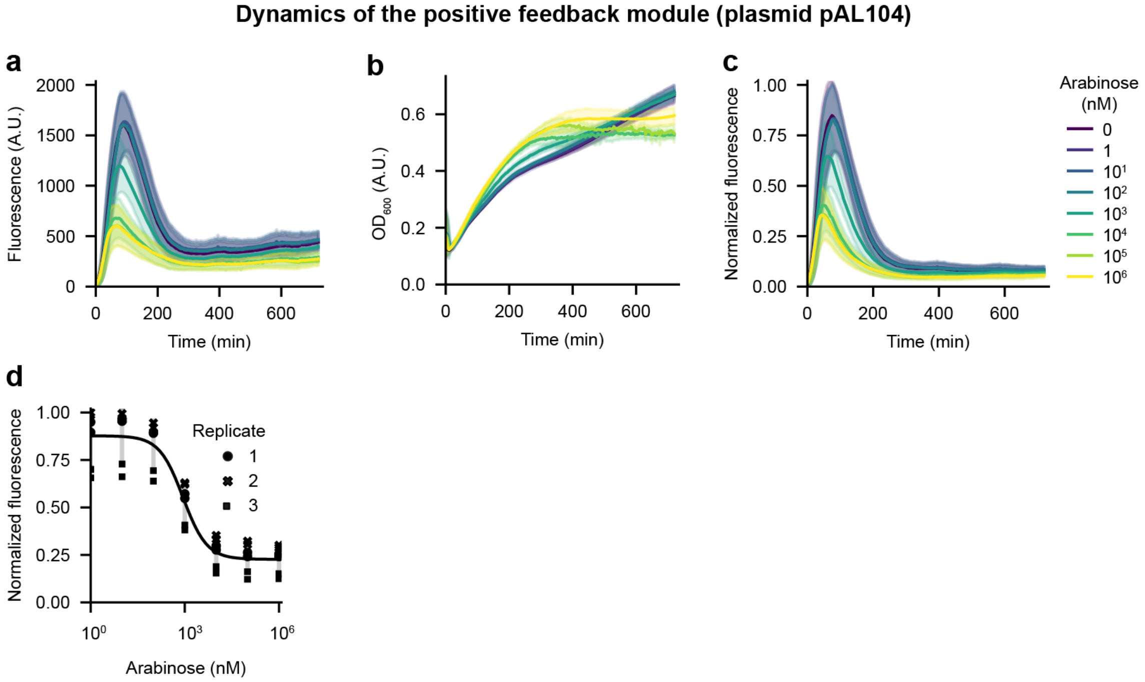

**Supplementary Figure 5 | Characterization of the positive feedback module (plasmid pAL104). (a-c)** Time-series plots showing sfGFP fluorescence (a), $OD_{600}$ (b) and normalized fluorescence/$OD_{600}$ (c) for the positive feedback strain carrying plasmid pAL104 with different arabinose treatments. n = 3 biological replicates with 2 technical replicates each. Error envelopes indicate standard deviation. **(d)** Dose-response curve at 3 h after treatment with arabinose for the data shown in (a-c). Error bars indicate standard deviation.

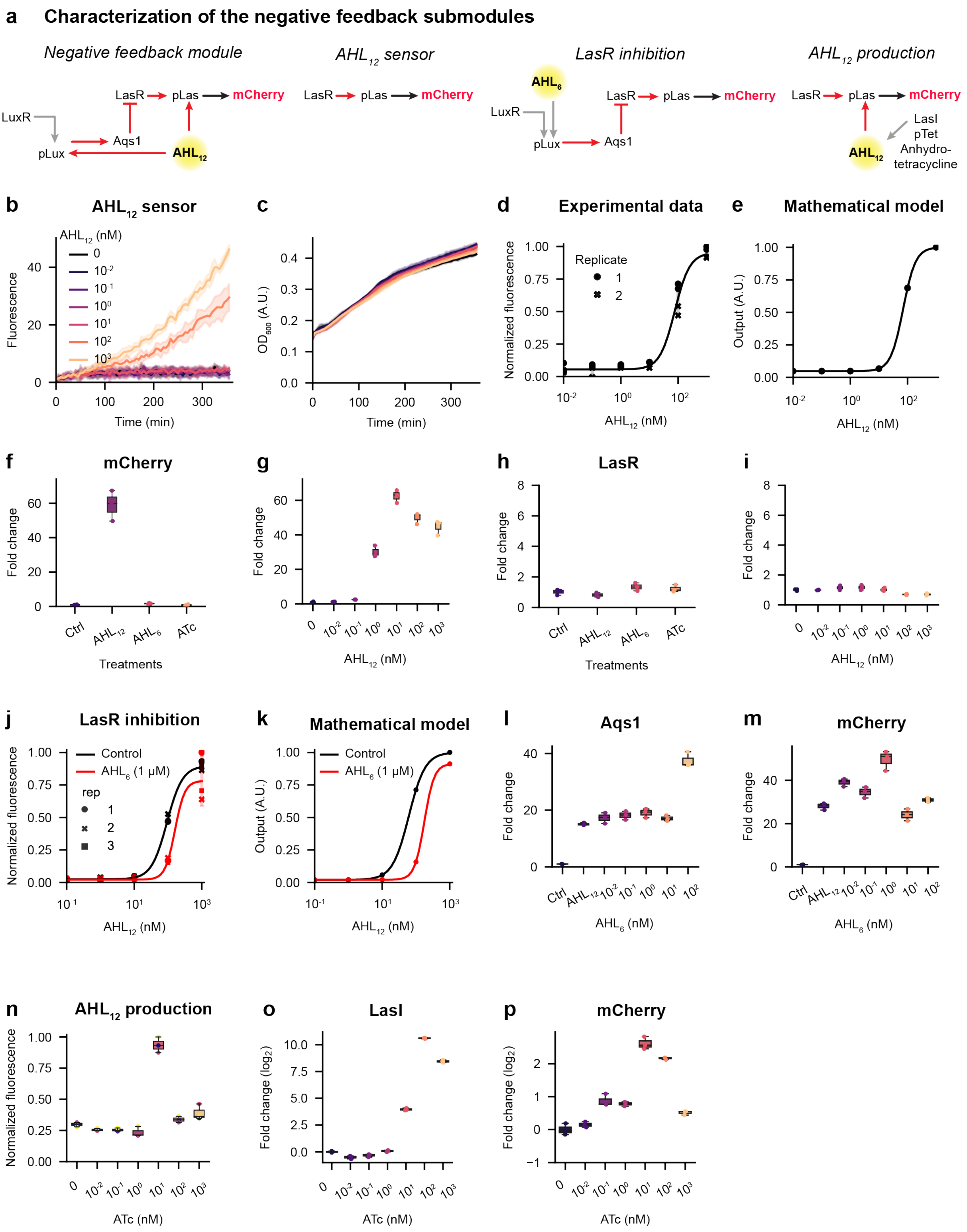

**Supplementary Figure 6 | Characterization of the negative feedback submodules. (a)** Schematics of the negative feedback module and submodules: $AHL_{12}$ sensor, $AHL_{12}$ sensor + LasR inhibition, and $AHL_{12}$ sensor + LasI production. **(b,c)** Time-series plots for the $AHL_{12}$ sensor strain showing mCherry fluorescence (b) and $OD_{600}$ (c) upon different $AHL_{12}$ treatments. n = 2 biological replicates with 2 technical replicates each, from the same experiment as in Fig. 2g. Error envelopes indicate standard deviation. **(d,e)** Experimental and simulated dose-response curves for the $AHL_{12}$ sensor strain showing normalized mCherry fluorescence upon treatment with indicated $AHL_{12}$ doses for 6 h. n = 2 biological replicates with 2 technical replicates each, from the same experiment as in Fig. 2g. (e) shows model simulation results. **(f-i)** RT-qPCR analysis for the $AHL_{12}$ sensor strain showing change in mRNA levels of *mCherry* and *LasR* after treatments with $AHL_{12}$ (1000 nM), $AHL_6$ (1000 nM), and ATc (1000 nM) in panels (f,h), and the indicated amounts of $AHL_{12}$ in panels (g,i) for 4 h. n = 1 biological replicate with 3 technical replicates. **(j,k)** Dose-response curves for the LasR inhibition strain at 6 h time-points for the data in Fig. 2i. The $AHL_{12}$ threshold concentration increases from 93 ± 2 nM to 160 ± 22 nM. (k) The mathematical model recapitulates these experimental findings. **(l,m)** RT-qPCR analysis for the LasR inhibition strain showing change in mRNA levels of (l) *Aqs1* and (m) *mCherry* upon treatment with different $AHL_6$ concentrations for 4 h. Note that all samples except the control were also treated with 1 µM $AHL_{12}$. n = 1 biological replicate with 3 technical replicates. **(n-p)** For the $AHL_{12}$ production strain, normalized mCherry output upon treatment with indicated ATc doses for 6 h (n), and RT-qPCR analyses (o,p) of change in *LasI* and *mCherry* mRNA levels of upon treatment with the indicated ATc concentrations for 4 h are shown. n = 1 biological replicate with 3 technical replicates. LasI: $AHL_{12}$ synthase, ATc: Anhydrotetracycline. *ihfβ* was used as a reference gene for all RT-qPCR analyses. In (d) and (j), error bars indicate standard deviation. In the boxplots, the center line represents the median, while the upper and lower box edges correspond to the first and third quartiles, respectively. Whiskers extend to the farthest data point within 1.5 times the interquartile range from the box edges or to the maximum data point within this range, whichever is closer.

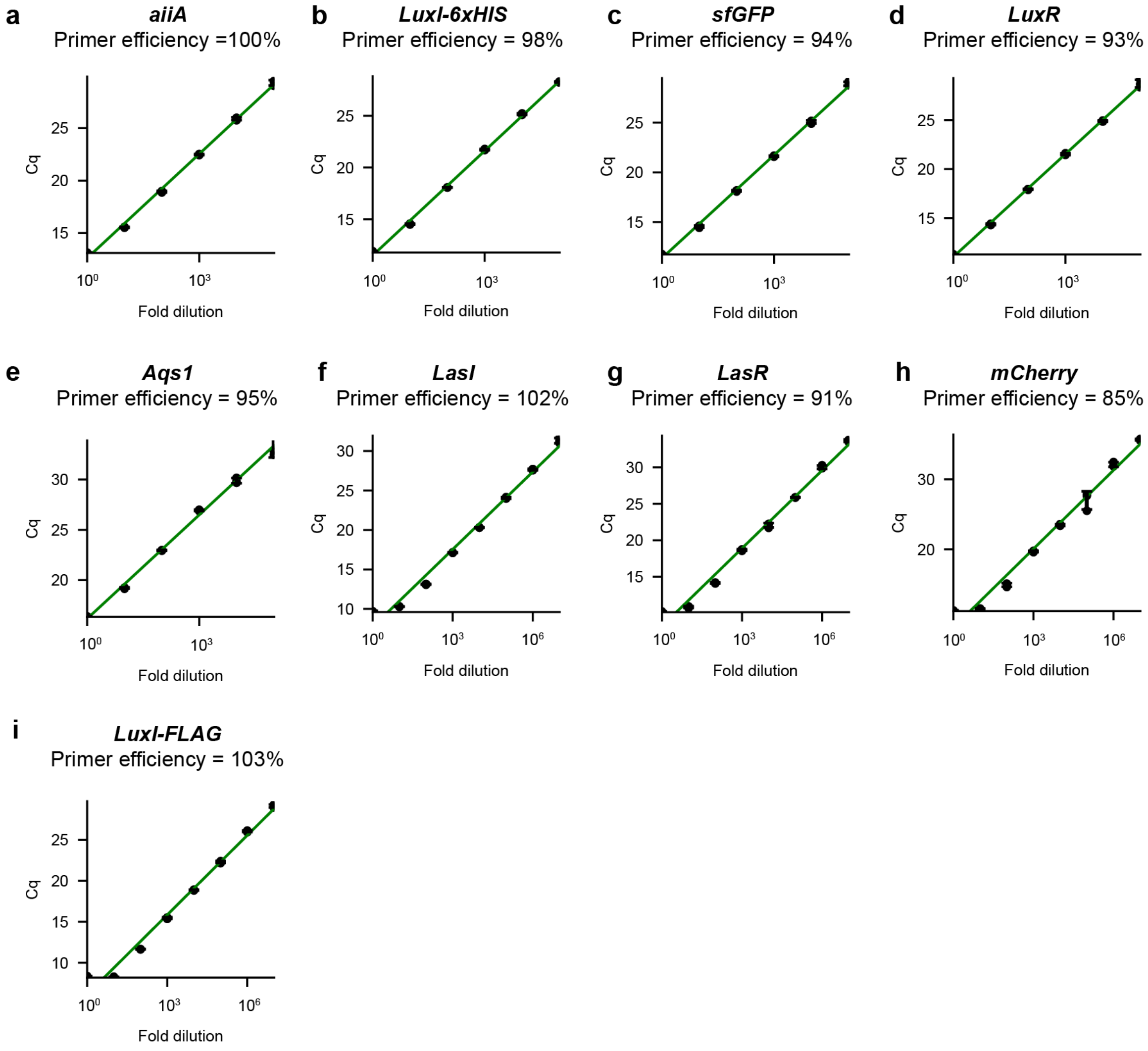

**Supplementary Figure 7 | qRT-PCR primer efficiency and standard curves. (a-i)** Standard curves showing cycle threshold (Cq) values for qPCR using the designated primer and serial dilutions of an appropriate template DNA. Experimentally determined primer efficiency values are shown. n = 1 biological replicate each with 3 technical replicates. Error bars indicate standard deviation.

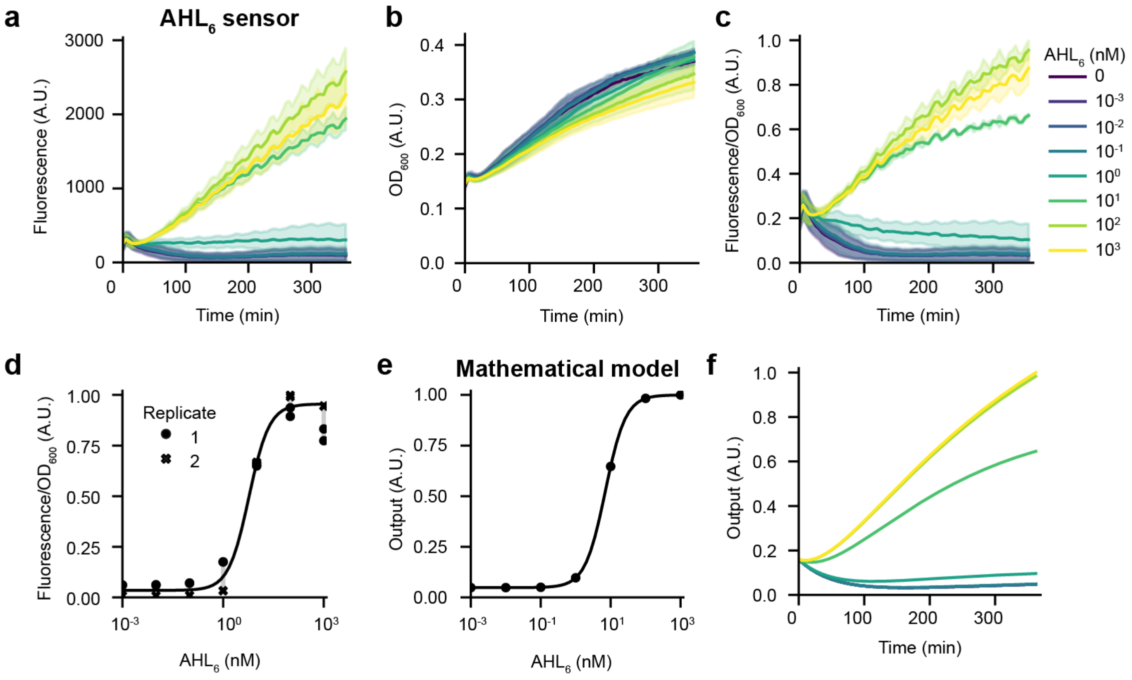

**Supplementary Figure 8 | $AHL_6$ sensor strain characterization and mathematical modeling. (a-c)** Time-series plots showing GFP fluorescence (a), $OD_{600}$ (b), and fluorescence/$OD_{600}$ (c) (scaled to 1 by dividing by the maximum) for the $AHL_6$ sensor strain upon different $AHL_6$ treatments. n = 2 biological replicates with 2 technical replicates each. Error envelopes indicate standard deviation. **(d)** Dose-response curve after 6 h $AHL_6$ treatment for the data shown in (a-c). A sigmoid function was fitted to the data to determine the activation threshold (6 ± 1 nM). Error bars indicate standard deviation. **(e,f)** Mathematical modeling of the $AHL_6$ sensor recapitulates the dose-response profile (e) as well as dynamics of fluorescence output (f).

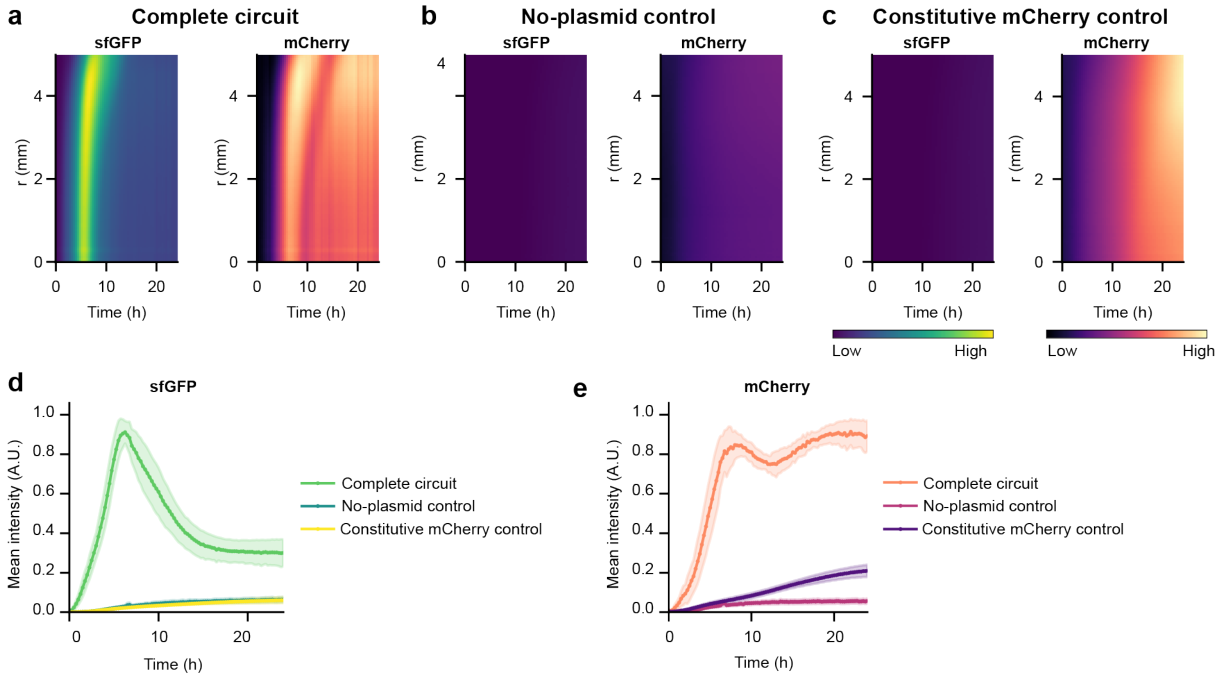

**Supplementary Figure 9 | Traveling waves are observed specifically for the complete circuit. (a-c)** Representative space-time plots of radial profiles of sfGFP and mCherry fluorescence for the complete circuit (a), no-plasmid control (b), and constitutive mCherry expression (c) strains. Data in (a) is the same as in Fig 4c,d. For (b) and (c), n = 3 biological replicates. **(d,e)** Mean intensities of sfGFP (d) and mCherry (e) for the indicated strains. The values are normalized to scale the maximum to 1. Error envelopes indicate standard deviation. Data for the complete circuit strain are the same as in Fig. 4.

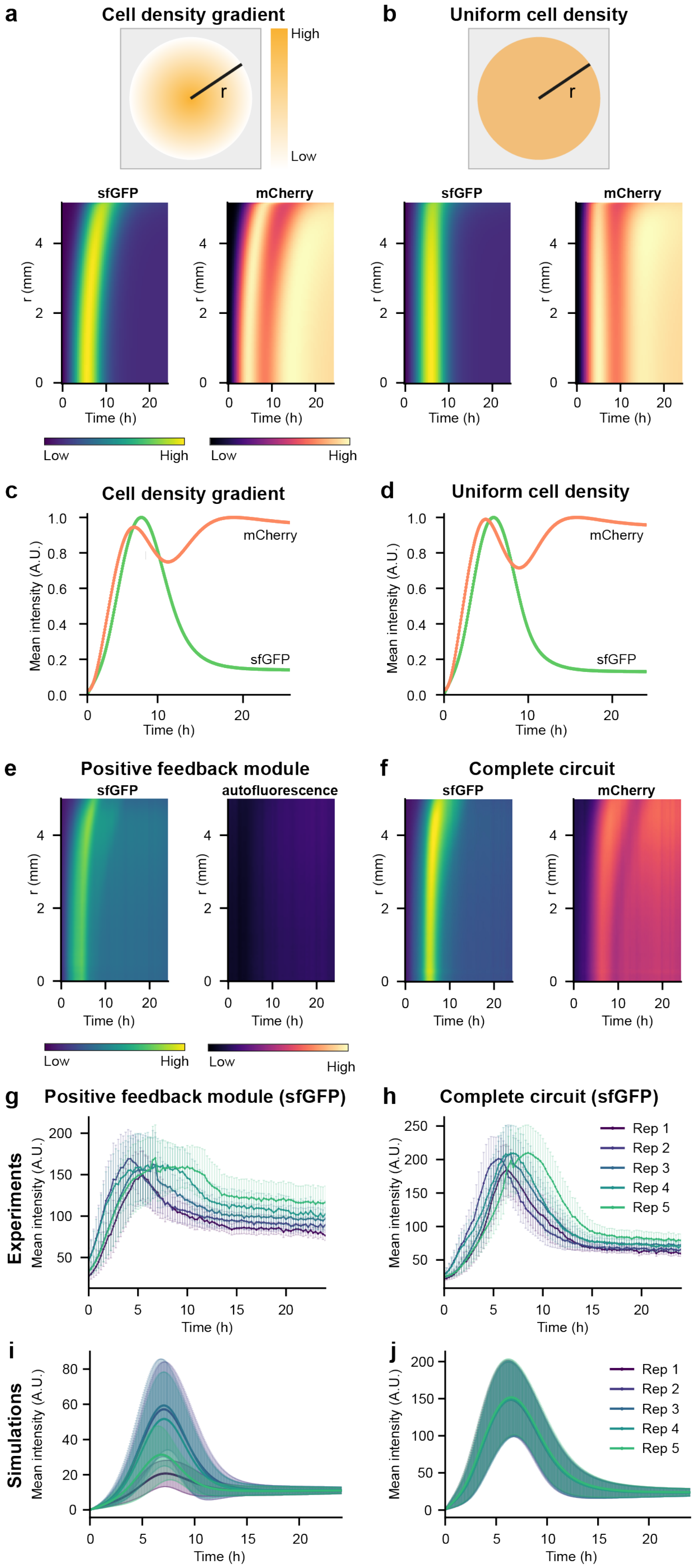

**Supplementary Figure 10 | Mathematical simulations reveal the role of cell-density gradients and feedback for traveling waves. (a,b)** Schematic showing the simulation setup along with space-time plots of simulated sfGFP and mCherry levels for graded cell density (a) and uniform cell density (b) conditions. **(c,d)** Mean sfGFP and mCherry intensities in the simulated cell lawn for the indicated cell density conditions. **(e,f)** Representative space-time plots of radial profiles of sfGFP and mCherry fluorescence for the positive feedback module (e) and the complete circuit (f). **(g,h)** Mean intensities of sfGFP fluorescence over time for the positive feedback module (g) and the full circuit (h), n = 5 biological replicates. **(i,j)** Simulation results showing mean intensities of sfGFP fluorescence over time for the positive feedback module (i) and the complete circuit (j), n = 5 stochastic simulations with noise terms. Error bars indicate standard deviation in (g-j).

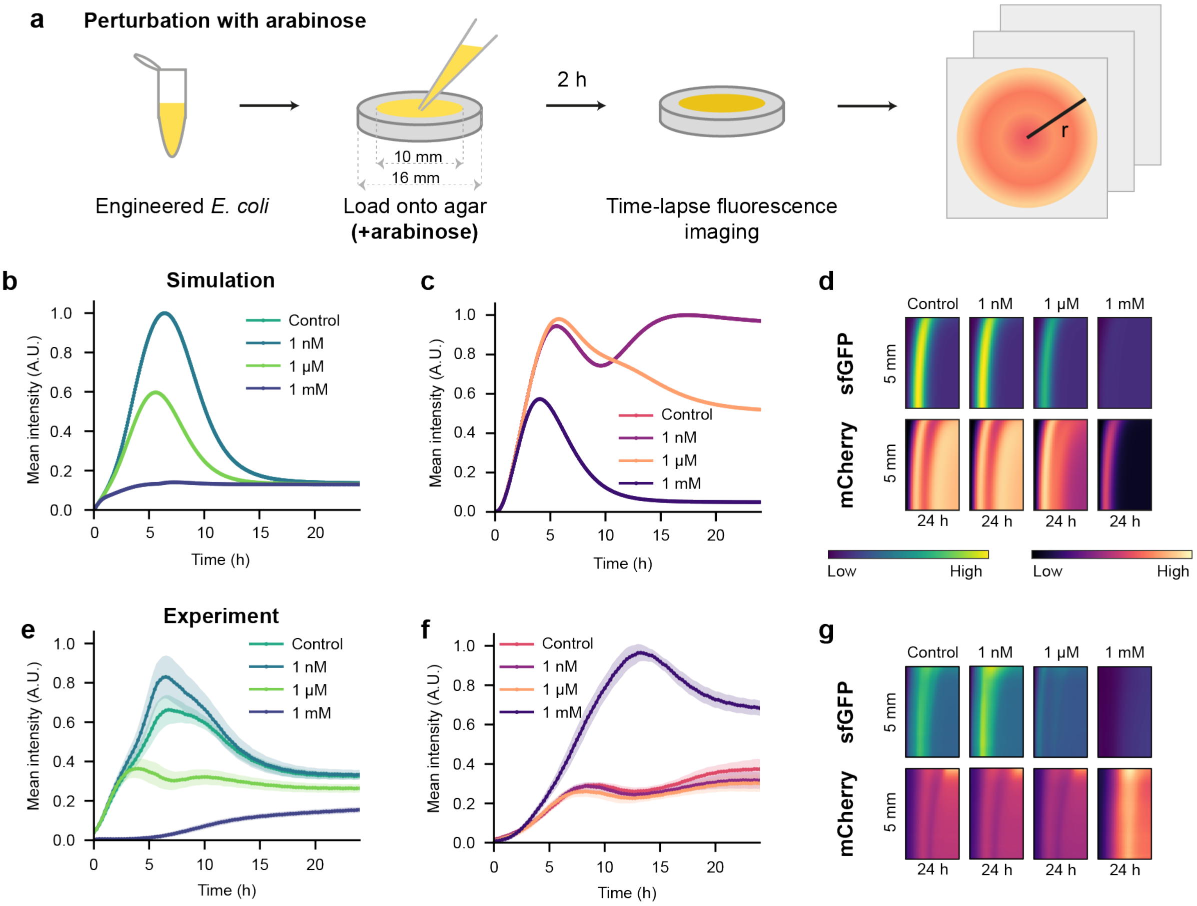

**Supplementary Figure 11 | Perturbation analysis with arabinose-induced AHL degradation.** **(a)** Experimental setup. **(b,c)** Simulated (b) sfGFP and (c) mCherry temporal dynamics for the indicated arabinose treatments. **(d)** Simulated space-time plots of sfGFP and mCherry for the indicated arabinose treatments. The simulations were performed by solving a PDE-based model (Model 5, Supplementary Note). **(e,f)** Mean intensities over time of sfGFP (e) and mCherry (f) for the indicated arabinose treatment conditions. The raw values were normalized by dividing through the maximum value of the dataset. Error envelopes indicate standard deviation. **(g)** Space-time plots of sfGFP and mCherry fluorescence for the indicated arabinose treatment conditions. The time-lapse imaging was performed for 24 h with 15 min intervals. n = 4 biological replicates.

### Supplementary Movie Legends

For all supplementary movies, *viridis* and *magma* lookup tables were used for sfGFP and mCherry, respectively. All images within the same time-lapse have the same gray value range. Supplementary Movie 3, Supplementary Movie 4, and Supplementary Movie 6 show results of 2D simulations.

**Supplementary Movie 1 | Complete circuit strain showing traveling waves.** The movie was recorded for 24 h with 15 min intervals. Traveling waves of fluorescence signal can be observed. Scale bar = 2 mm. n = 9 biological replicates.

**Supplementary Movie 2 | Control strains do not show traveling waves.** The movie was recorded for 24 h with 15 min intervals. The no-plasmid strain shows low levels of autofluorescence in GFP and mCherry channels. The constitutive mCherry expression strain shows autofluorescence in the GFP channel and elevated levels of fluorescence signal in the mCherry channel. Scale bar = 2 mm. n = 6 biological replicates.

**Supplementary Movie 3 | Simulation of traveling waves with graded and uniform initial cell density.** Top: Graded cell-density, bottom: uniform cell-density, left: sfGFP, right: mCherry.

**Supplementary Movie 4 | Simulation of traveling waves with different lawn sizes.** Left: sfGFP, right: mCherry.

**Supplementary Movie 5 | Effect of lawn size on traveling wave patterns.** The movie was recorded for 24 h with 15 min intervals. Three lawn sizes are shown. Scale bar = 2 mm. n = 4 biological replicates.

**Supplementary Movie 6 | Simulation of traveling waves with different lawn positions.** Left: sfGFP, right: mCherry.

**Supplementary Movie 7 | Effect of lawn position on traveling wave patterns.** The movie was recorded for 24 h with 15 min intervals. Three lawn positions are shown. Scale bar = 2 mm. n = 6 biological replicates.

**Supplementary Movie 8 | Effect of arabinose-induced AHL degradation on traveling waves.** The movie was recorded for 24 h with 15 min intervals. Four arabinose treatments are shown. Scale bar = 2 mm. n = 1 biological replicate.
